# Parthenolide boosts megakaryocyte maturation and restores platelet responses in Wiskott–Aldrich syndrome

**DOI:** 10.64898/2026.09.13.751254

**Authors:** Rhaissa Calixto Vieira, Lia Goncalves Pinho, Tracer Yong, Minghui He, Giulio Zanette, Marta Dominguez Gargallo, Elisabet Jene Vinuesa, Isabella Rosales Enriquez, Olov Ekwall, Anna-Lena Gustavsson, Lisa S. Westerberg

## Abstract

Wiskott-Aldrich syndrome (WAS) is an inborn error of immunity with a broad disease spectrum, classified into class I (late onset) or II (early onset) variants. Thrombocytopenia and small platelets are the most consistent findings among patients, difficult to treat and related to development of autoimmunity. To identify new treatment options for thrombocytopenia in WAS, we developed a FACS-based screening for drug repurposing. We identified parthenolide as a lead small molecule that increased WASp abundance in cells with residual WASp expression. Using the megakaryocytic MEG-01 cells, gene edited to express WAS class I and II genetic variants, parthenolide induced megakaryocyte maturation as evident by upregulation of CD61, increased cell size and complexity, increased phosphorylation of ERK1/2, and higher DNA ploidy. We generated a new mouse model harboring a WAS class I missense variant WASp-R88C, corresponding to human WASp-R86C, with reduced expression of WASp. WASp-R88C mice had lower numbers of platelets compared to WT mice. Bone marrow-derived WASp-R88C and WASp-KO megakaryocytes, differentiated in the presence of parthenolide derivative, DMAPT, showed increased ploidy and upregulation of the maturation markers CD61, CD41 and CD42d, resulting in improved platelet production *in vitro*. Treatment with parthenolide derivative, DMAPT, led to increased platelet numbers *in vivo* in WASp-R88C mice and dampened the hyperactivation of WAS patient platelets by reducing thrombin-induced CD62P exposure after activation. The identification of parthenolide offers a promising therapeutic approach for WAS patients who are unresponsive or unsuitable for definitive therapies.

**Key points:**

- A FACS-based drug screen using degradation-prone class I WAS variants identifies parthenolide/DMAPT as a candidate that enhances megakaryocyte maturation and platelet production *in vitro*.
- Parthenolide/DMAPT thrombopoietic activity increases platelet production *in vivo* and restores improves cytoskeletal remodeling of WAS megakaryocytes and platelets.

## Introduction

Wiskott–Aldrich syndrome (WAS) is an inborn error of immunity caused by genetic variants in the *WAS* gene encoding the hematopoietic-specific actin regulator WAS protein (WASp) (1,2). Patients have a spectrum of clinical manifestations, including bleeding, recurrent infections, autoimmunity, and malignancy (3). Thrombocytopenia with small platelets is a consistent and early feature of both late onset (class I) and early onset (class II) disease and is attributed to defects in megakaryocyte development, platelet function and the presence of autoantibodies (4–10).

Disease severity correlates with the level of WASp expression. Class II variants typically result in a complete absence of WASp, whereas class I variants with missense mutations in exons 1 and 2, preserve residual protein expression but lead to reduced WASp levels in hematopoietic cells (11–14). In class I WAS, WASp mRNA is normally produced but the mutant protein undergoes accelerated degradation, in part due to impaired interaction with the WASp-interacting protein (WIP) that is a key stabilizing partner for WASp (15). This post-translational instability identifies class I WAS as a particularly relevant setting in which increasing WASp abundance may restore cellular function. Supporting this concept, a WIP-derived peptide has been shown to stabilize mutant WASp and partially rescue immune cell phenotypes *in vitro*, although this strategy has not progressed to clinical application (16).

Current treatment options include hematopoietic stem cell transplantation and gene therapy, which are curative but not universally accessible or suitable (17–19). Patients with milder disease are often managed with thrombopoietin receptor agonists, such as eltrombopag or romiplostim (20). However, responses are variable and platelet defects are not fully corrected (21–23). These limitations highlight a continued need for therapeutic strategies that directly address the underlying cellular dysfunction in WAS, particularly in patients with residual but unstable WASp expression.

The lack of targeted therapies for thrombocytopenia in WAS underscores the need for new therapeutic approaches. Drug repurposing offers an efficient strategy to identify compounds capable of restoring cellular function in disease-relevant models (24). Using a high-throughput FACS based drug screening approach, we identified parthenolide that promotes megakaryocyte maturation and platelet responsiveness in WAS murine and human cells, uncovering a potential therapeutic approach for inherited thrombocytopenia characterized by impaired platelet production and function.

## Methods

### Human samples

Washed platelets from healthy donors were obtained from buffy coats of anonymous blood donors processed by Karolinska Universitetssjukhuset and upon written consent. Blood from patients and family caregivers was provided upon written consent. Blood was collected in citrate tubes and received upon 24h after withdrawing. The Code of Ethics of the World Medical Association (Declaration of Helsinki) for human samples was followed; the study was approved by the institutional Ethical Committee (EPM# 2023-04773).

### Generation of Was-R88C mice

To generate a mouse model carrying the class I WAS-associated variant corresponding to human WASp-R86C, a knock-in point mutation was introduced into the murine *Was* locus using CRISPR-Cas9 genome editing. The murine R88C substitution corresponds to the human WASp-R86C variant. CRISPR-Cas9 targeting was performed in collaboration with Karolinska Genome Engineering and the Karolinska transgenic animal core facility. C57BL/6 wildtype, WASp knockout (WASp-KO), and WAS-R88C mice were bred under specific-pathogen-free conditions in the animal facility of KM Wallenberg, Karolinska Institutet. Mice aged 6–10 weeks were used for experimental treatments. All animal experiments performed were approved by the Stockholm North Animal Ethics Committee permit #05294-2023.

### Generation of WAS knockout (WKO) MEG-01 cells

*WAS* KO MEG-01 cells were generated by CRISPR/Cas9. Four different gRNA sequences targeting *WAS* were generated and cloned into lentiGuide-Puro-P2A-EGFP-mRFPstuff (Addgene #137730). For cloning, the 25bp oligos (Table 1; ordered from Thermo Fisher Scientific) were annealed and phosphorylated with T4 PNK (NEB, 0201S) and subsequently used in a ligation reaction with Bsmb1-digested lentiGuide-Puro-P2A-EGFP_mRFPstuf vector using Quick Ligase (NEB, M2200S). Transformed One Shot™ Stbl3 cells (Thermo Fisher, C737303) were plated on LB-ampicillin agar and incubated overnight at 37 °C. Individual colonies were inoculated into LB-ampicillin broth, grown with shaking at 37 °C for 16 h, and stored as 25% glycerol stocks. Plasmid DNA from each gRNA clone was purified using the QIAprep® Spin Miniprep Kit (Qiagen) and verified by Sanger sequencing.

**Table 1:**
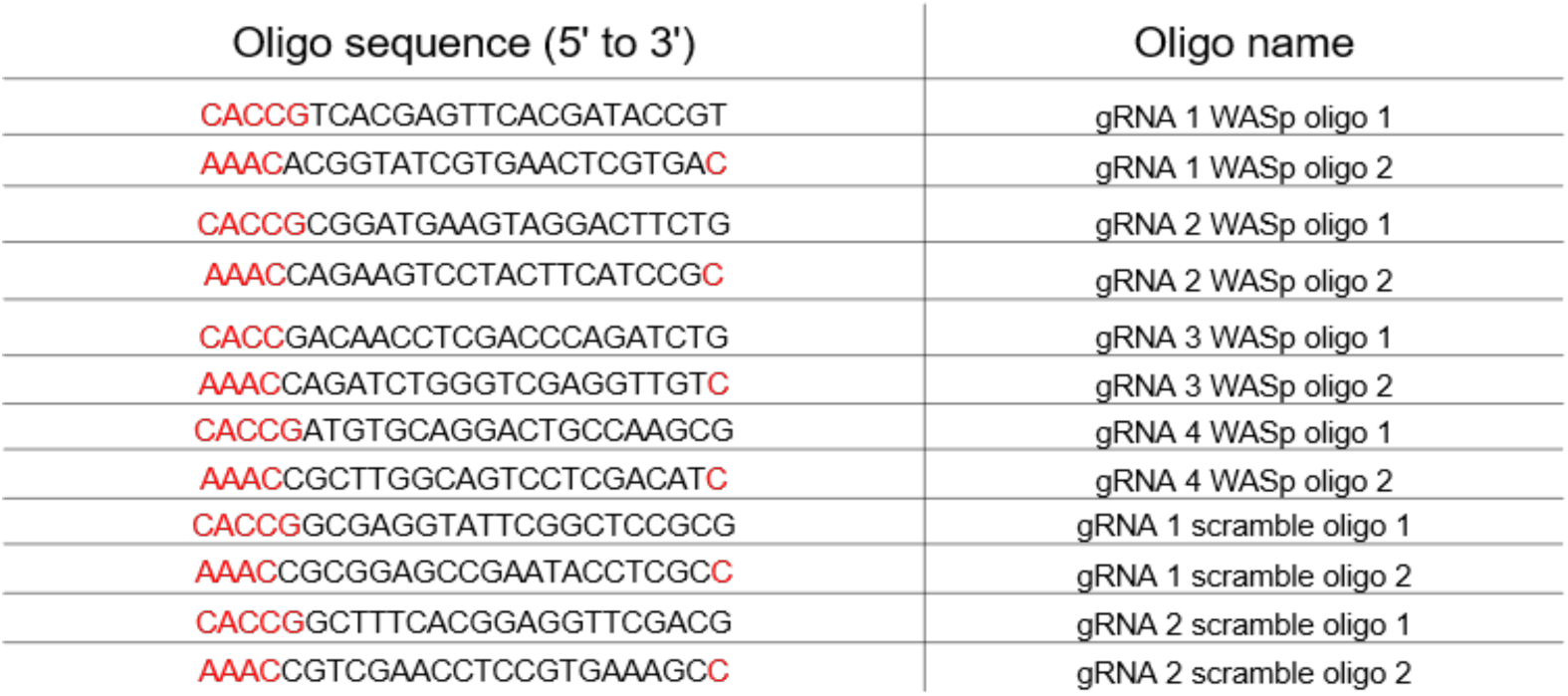
Oligonucleotide sequences used for generation of CRISPR-Cas9 gRNA constructs targeting the WASp gene and scramble control sequences. Sequences in red indicate cloning overhangs (CACC and AAAC) used for directional cloning into expression vectors via BbsI restriction enzyme sites.

MEG-01 cells were lentivirally transduced to express Cas9-BFP (Addgene #78545). Sorted BFP-positive cells were electroporated (Amaxa 4D, FF-120, homemade buffer) with lentiGuide-Puro-P2A-EGFP gRNA-cloned. After 48h, EGFP/BFP double-positive cells were sorted by flow cytometry. Mismatch cleavage assay was used for polyclonal screening to determine which gRNA that provided most detectable NHEJ (Surveyor® Mutation Detection Kit - S100, IDT). Cells targeted with gRNA1 and gRNA2 were clonally expanded. Clones with absent or low WASp levels were further sequenced to identify indels spanning 300bp upstream and downstream of the Cas9-targeted region. WASp KO cells were confirmed by WB.

### Compound Library

Screening volumes of a total of 7172 compounds were obtained from the Chemical Biology Consortium Sweden, SciLifeLab including FDA/EMA approved.

### Flow Cytometry-Based drug screening

To develop the high-throughput screening assay, MEG-01 cells expressing EGFP-WASp-R86C were used. The compound library (stock solutions of 10 mM DMSO solutions) was pre-spotted by an automated liquid handling system. were pre-spotted by an automated liquid handling system. The WASp-R86C cells were seeded onto pre-spotted molecules (final concentration of 10 µM) in 96-well plates containing cycloheximide (CHX) and incubated overnight. Compounds resulting in an accumulation index (AI) greater than 1.2 in the primary screening were selected as positive hits. Selected positive hits underwent secondary validation using antibody-based detection of WASp (anti-WASp conjugated to mouse-AF647), with molecules tested at 3 concentrations. Molecules that maintained consistent WASp stabilization across multiple detection methods (EGFP and antibody staining) were confirmed as validated hits.

## Results

### Class I WAS variants exhibit reduced WASp stability

To model the degradation characteristics of WASp caused by missense variants and directly assess their impact on megakaryocytes and platelets, we used CRISPR-Cas9 to gene edit the *WAS* gene in the megakaryocytic cell line MEG-01 (Figure 1A). We identified WAS knockout (KO) MEG-01 cells (Figure 1A–C). Moreover, we identified clone #1 from the gRNA2-targeted cells with reduced WASp expression (Supplementary Figure 1A). Sequencing analysis indicated a lysine (K) codon insertion between K81_S82 (Supplementary Figure 1B), hereafter referred to as WASp-insK82. This region is within the WASp WH1 domain for interaction with WIP (WASp aa 34-149), where WIP masks residues K76 and K81 and inhibits their ubiquitylation (15,26). The insertion of another lysine next to K81 may have impacted WIP-WASP interaction or ubiquitylation resulting in reduced WASp expression (Figure 1B, C).

**Figure 1:**
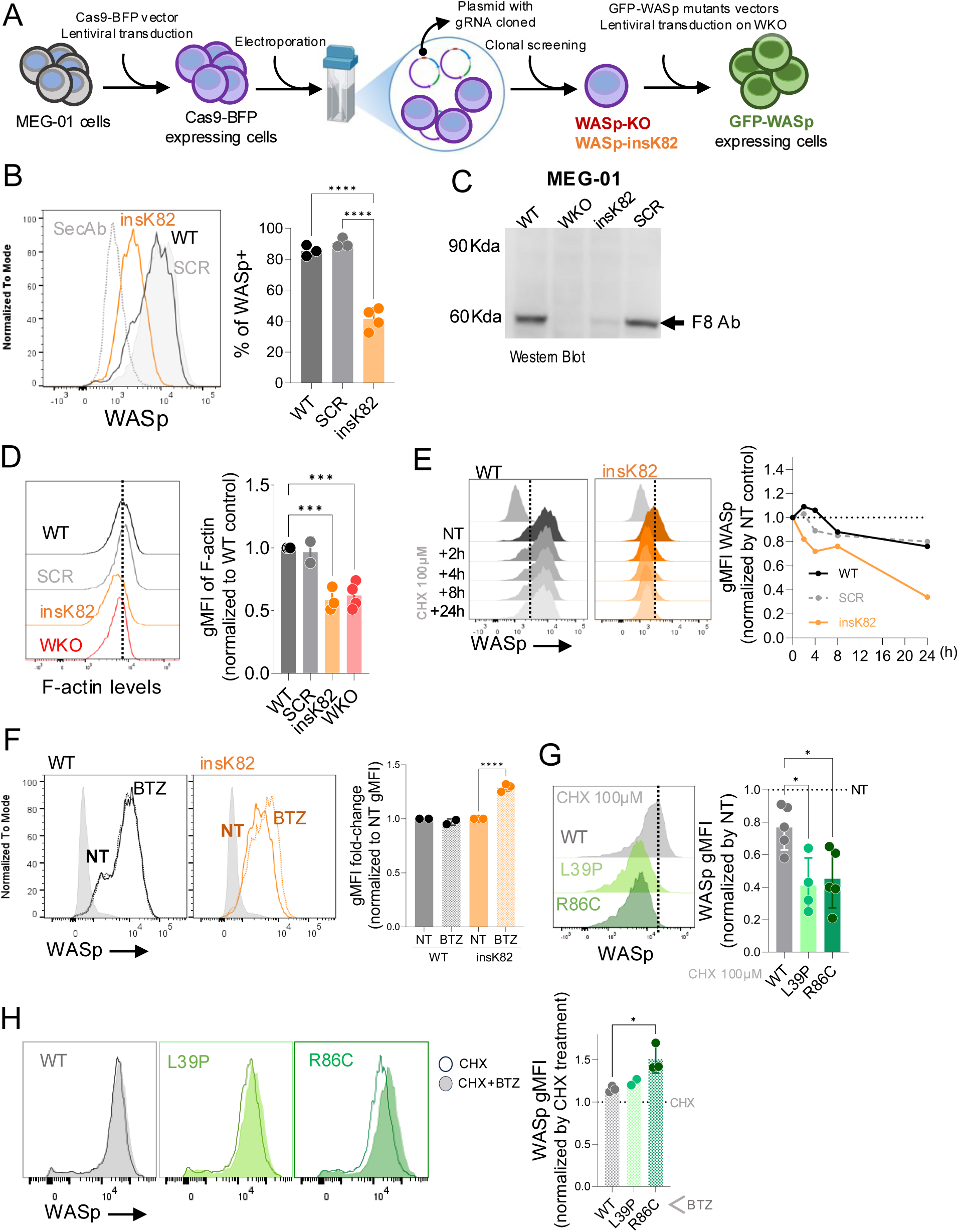
Generation of genetically engineered megakaryocyte cell line models to detect WASp degradation. **(A)** Scheme summarizing the pipeline for generation of different MEG-01 cells. WASp expression in MEG-01 cells lines detected by **(B)** Flow cytometry and **(C)** Western Blotting. **(D)** F-actin levels in MEG-01 cells. Left: representative histogram. Dotted line marks the mean value in WT samples. Right: quantification of gMFI values relative to WT samples. **(E)** WASp-chase assay after cycloheximide (CHX) treatment. Left: representative histograms of WASp expression over time. Right: quantification of gMFI values normalized to non-treated (NT) controls. Vertical dotted line indicates the point of positive and negative staining for WASp based on the secondary antibody background. **(F)** WASp detection after 6h of proteasome inhibition by Bortezomib (BTZ) 1µM. Left: representative histograms with NT and BTZ-treated samples; right: quantification of gMFI values normalized by NT cell specific samples. **(G)** MEG-01-engineered cells in WASp-chase assay after CHX treatment overnight. Left: Representative histograms; dotted line shows mean values of WT samples. Right: WASp gMFI normalized by cell specific NT control; values below 1 show a mean of 20% of WASp-WT degradation, and 60% for WASp-L39P and WASP-R86C. **(H)** WASp detection in samples treated overnight with CHX or CHX+BTZ. Quantification shows WASp gMFI values normalized by cell specific CHX control; values above 1 indicate that degradation is prevented. One-way ANOVA followed by (A and F) Tukey’s multiple comparisons test; (B, G and H) Dunnett’s multiple comparisons test. Experiments have at least n = 3 experimental replicates. Each dot represents one biological experiment.

WASp-KO and WASp-insK82 cells displayed reduced F-actin content, indicating impaired cytoskeletal regulation (Figure 1D). To directly assess protein stability, we performed cycloheximide (CHX) chase experiments to block de novo protein synthesis. In WASp-insK82 cells, WASp levels declined rapidly and were nearly undetectable after 24 hours, in contrast to the relatively stable expression observed in WT cells (Figure 1E). Inhibition of the proteasome with bortezomib (BTZ) partially rescued WASp levels in WASp-insK82 cells, supporting a degradation mechanism involving proteasomal pathways (Figure 1F).

To model clinically relevant class I variants, we expressed EGFP-tagged WASp constructs carrying the L39P or R86C mutations in WASp-KO MEG-01 cells (Supplementary Figure 1C). While baseline expression levels were comparable across constructs, CHX treatment revealed reduced stability of EGFP-WASp-L39P and EGFP-WASp-R86C compared to EGFP-WASp-WT (Figure 1G and Supplementary Figure 2A-B). Proteasome inhibition led to accumulation of mutant WASp, particularly for WASp-R86C, whereas WASp-WT remained largely unaffected (Figure 1H), supporting enhanced proteasomal turnover of class I WAS variant proteins.

Together, these models recapitulated key features of class I WAS, characterized by reduced WASp stability with increased proteasomal degradation. These findings established a disease-relevant platform to identify small molecules that increase WASp abundance and restore cytoskeletal function.

### Drug repurposing screening identifies parthenolide as a lead candidate

To identify compounds capable of increasing WASp abundance in cells expressing unstable variants, we established a flow cytometry-based drug screening strategy using MEG-01 cells expressing the class I variant WASp-R86C. Assay development and optimization demonstrated robust performance (Z-score>0.5) under conditions of inhibited protein synthesis and was used for compound screening (Supplementary Figure 3).

We screened the Prestwick FDA/EMA-approved library (1,260 compounds) and the SPECS Drug Repurposing library (∼6,000 compounds) using as read-out the EGFP(-WASp) signal (Figure 2A, B; Supplementary Figure 4A, B). The screening assay showed consistent performance across plates, with Z-factors above 0.5 (Figure 2C). Primary screening identified 27 candidate compounds from the Prestwick library and 60 candidate compounds from the SPECS library that increased the EGFP signal above the predefined threshold (Figure 2D and Supplementary Figure 4C, D).

**Figure 2:**
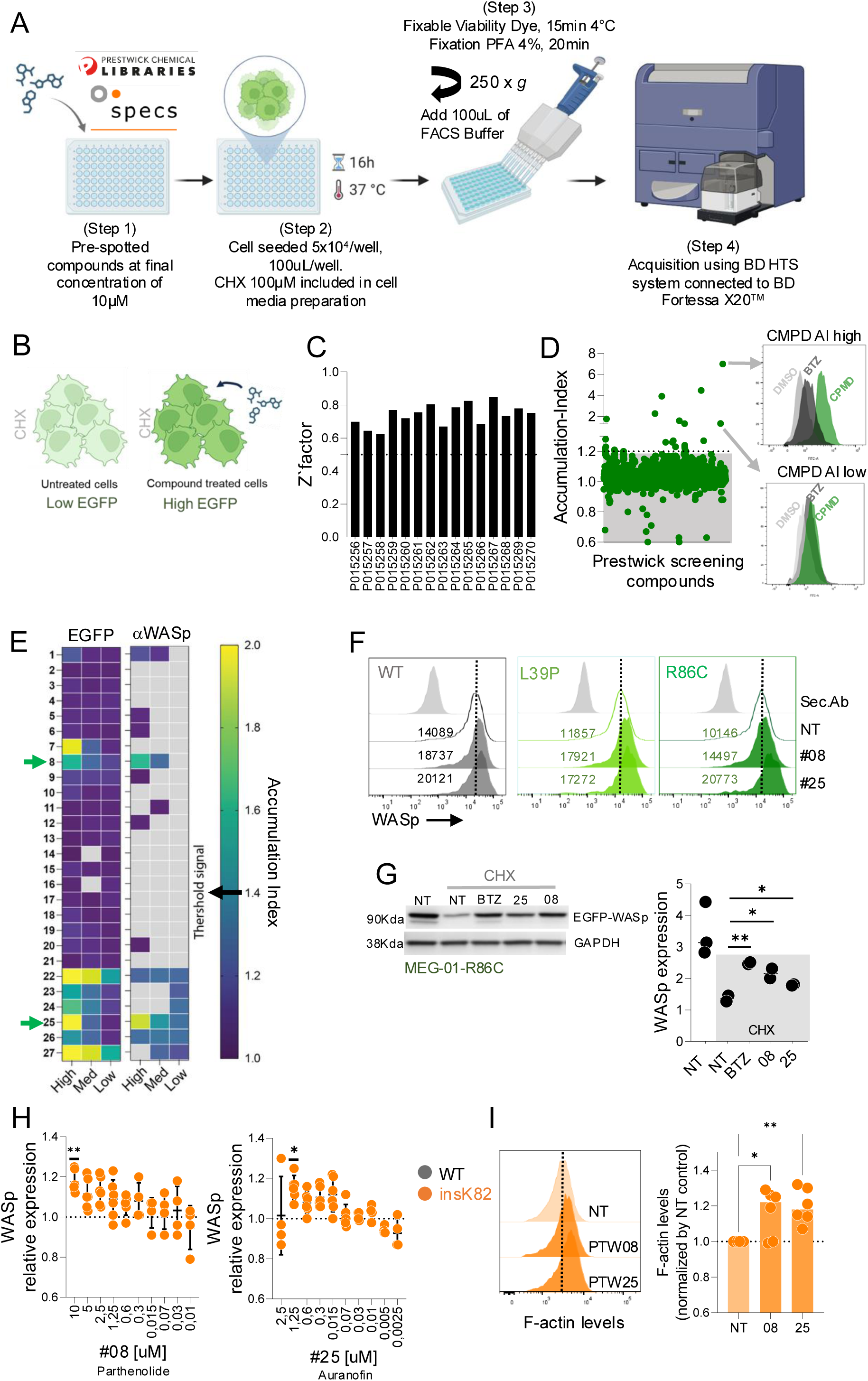
Drug repurposing screening identifies parthenolide as a lead candidate. **(A)** Scheme summarizing the pipeline for screening. **(B)** Rational of the screening assay: MEG-01-R86C cells were treated with cycloheximide (CHX) to inhibit de novo translation. In the absence of a stabilizing compound, EGFP-WASp is rapidly degraded, resulting in low EGFP signal. In contrast, treatment with compounds that prevent WASp degradation leads to higher EGFP accumulation, serving as the screening read-out. **(C)** Z-factor values of each plate from the Prestwick library screening. (**D)** EGFP accumulation-index of the Prestwick screened compounds; compounds selected as positive are above the 1.2 AI threshold. **(E)** Heat maps of WASp accumulation-index of the 27 compounds from the first screening; left: EGFP signal, right: WASp-AF647 signal; compounds selected as positive are above 1.4 Accumulation-Index threshold. High, Med, Low refers to the compounds concentrations pre-spotted on assay plates. The gray squares are the values bellow Accumulation-Index = 1. **(F)** Representative histograms of WASp staining in EGFP-expressing WASp WT, WASp-L39P, and WASp-R86C cells showing the reduced WASp degradation in cultures treated with #08 (parthenolide) or #24 (auranofin); the numbers in the histogram are the gMFI**. (G)** Western blotting (left) and graph quantification (right) of WASp-R86C lysates incubated or not with #08 (parthenolide) or #24 (auranofin); BTZ as positive control. GAPDH is used as loading control. Right: quantification of WASp protein levels normalized to NT-CHX condition; shaded region on graph indicates samples treated with CHX. **(H)** WASp level relative to NT culture in WASp-insK82 MEG-01 for the indicated drug concentrations; cells were treated for 48h. **(I)** Representative histograms (left) and quantification (right) of F-actin levels in MEG-01 WASp-insK82 cells treated at optimal concentrations (#08: 10 µM; #25: 1.25 µM). Vertical dotted line represents mean NT control. Statistical analysis: one-way ANOVA followed by (E, F) Šídák’s multiple comparisons test or (G, H) Dunnett’s multiple comparisons test. F-I: Experiments have at least n = 3 experimental replicates. Each dot represents one biological experiment.

To exclude false-positive candidates, hits were evaluated using an orthogonal validation strategy combining direct WASp detection with the EGFP signal. Among the Prestwick hits, #08 and #25 induced increased EGFP and WASp signals and were therefore selected for further analysis (Figure 2E). Increased WASp abundance was confirmed in cells expressing the class I variants L39P and R86C (Figure 2F and Supplementary Figure 3I). Western blot analysis verified accumulation of WASp, excluding autofluorescence-related artifacts (Figure 2G). To assess the compounds in a model expressing endogenous WASp, we next examined the WASp-insK82 cells (Figure 1D). Both #08 and #25 increased WASp levels and were associated with elevated F-actin content, indicating improved cytoskeletal responses (Figure 2G-H). Compound #08 was identified as parthenolide, a feverfew-derived sesquiterpene lactone previously reported to enhance platelet production from megakaryoblastic cell lines and primary megakaryocytes (27–29). Compound #25 was identified as auranofin, associated with adverse effects on platelet viability (28,29). Together, parthenolide was selected as the lead compound for downstream functional studies.

### Parthenolide enhances megakaryocyte maturation and thrombopoietic responses in WAS cellular models

Based on our aim to enhance platelet production in WAS, we examined if parthenolide could promote terminal megakaryocyte maturation that culminates with thrombopoietic output in WAS cellular models. WT, WASp-insK82, and WASp-KO MEG-01 cells were used as MEG-01 cells maintain their intrinsic ability to mature and release platelet-like particles upon stimulation (30) (Supplementary Figure 5A, B). We examined megakaryocyte maturation characterized by acquisition of lineage-specific markers, cellular enlargement, polyploidization, cytoskeletal remodeling, and ultimately platelet production (31,32). Parthenolide treatment of WT, WASp-insK82, and WASp-KO MEG-01 cells increased the proportion of CD41-positive cells and promoted polyploidization within the CD41-positive population, while CD41-negative cells remained unaffected (Figure 3A-B; Supplementary Figure 5C). Consistent with progression toward a more mature phenotype, parthenolide increased cellular size and complexity (Supplementary Figure 5D). These changes were accompanied by enhanced phosphorylation of ERK1/2 (Figure 3C), a signaling pathway implicated in megakaryocyte differentiation, polyploidization, and platelet production (33), supporting activation of a maturation-associated signaling response.

**Figure 3:**
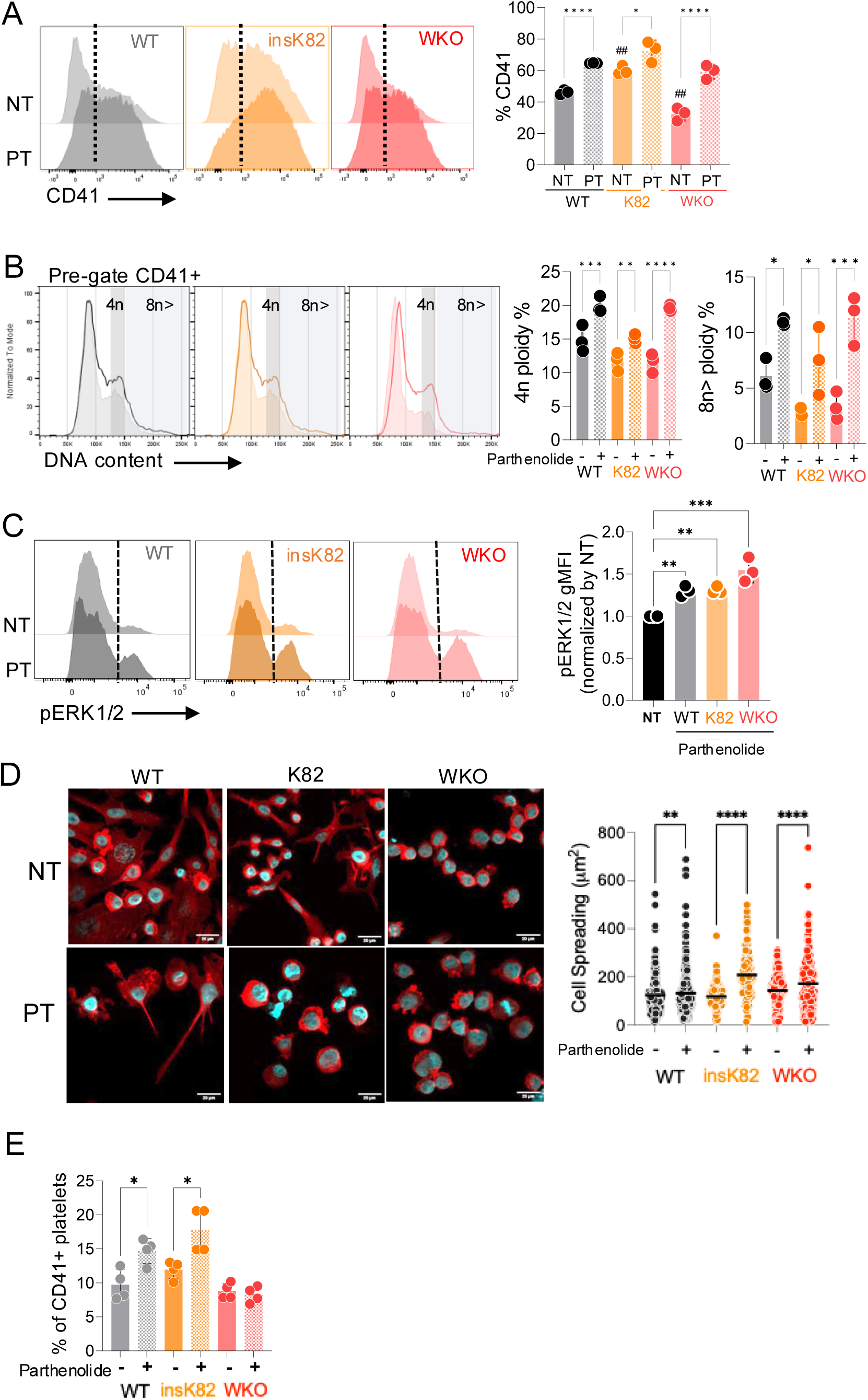
Parthenolide promotes maturation of MEG-01 cells. **(A)** CD41 expression in WASp-WT, WASp-insK82, and WASp-KO (WKO) MEG-01 cells treated or not (NT) with parthenolide (10µM, 48h). Left: representative histograms. Right: quantification of CD41⁺ cell percentages. **(B)** DNA content analysis (DAPI) of CD41⁺ gated MEG-01 cells. Left: representative histograms. Right: quantification of 4n and ≥8n ploidy across genotypes and treatment conditions. **(C)** Flow cytometry analysis of phosphorylated ERK1/2 (pERK1/2) expression in SCR, insK82, and WKO cells treated or not with parthenolide. Left: representative histograms. Right: quantification of pERK1/2 gMFI relative to NT controls. **(D)** Confocal images of WASp-WT, WASp-insK82, and WASp-KO MEG-01 cells seeded onto fibrinogen (100ug/mL) cultured overnight with or without parthenolide and stained for F-actin (phalloidin, red) and nuclei (DAPI, blue). Scale bar = 20 µm. Right: quantification of total cell spreading area. **(E)** Quantification of CD41⁺ platelet-like particles after 48h parthenolide treatment in WT, insK82, and WKO cultures. Statistical analysis: one-way ANOVA followed by Holm-Šídák’s multiple comparisons test (A–F). Experiments have at least n = 3 experimental replicates. Each dot represents one biological experiment.

Given the central role of the actin cytoskeleton in megakaryocyte maturation, we next examined cytoskeletal remodeling following adhesion to fibrinogen-coated surfaces. Parthenolide treatment enhanced cell spreading and cortical F-actin organization, indicating improved actin-dependent cellular responses (Figure 3D). Importantly, these phenotypic changes translated into enhanced thrombopoietic output, as parthenolide-treated cultures released higher numbers of platelet-like particles into the culture supernatant (Figure 3E).

To explore primary megakaryocyte maturation, we generated a mouse model expressing the WASp-R88C variant, corresponding to human hot spot variant WASp-R86C (13), and compared cellular responses of WT, WASp-R88C, and WASp-KO mice. Bone marrow-derived, expanded primary megakaryocytes from WT mice expressed WASp, while WASp-KO cells were devoid of WASp and WASp-R88C megakaryocytes had reduced WASp expression, modeling class II and class I WAS, respectively (Figure 4A).

**Figure 4:**
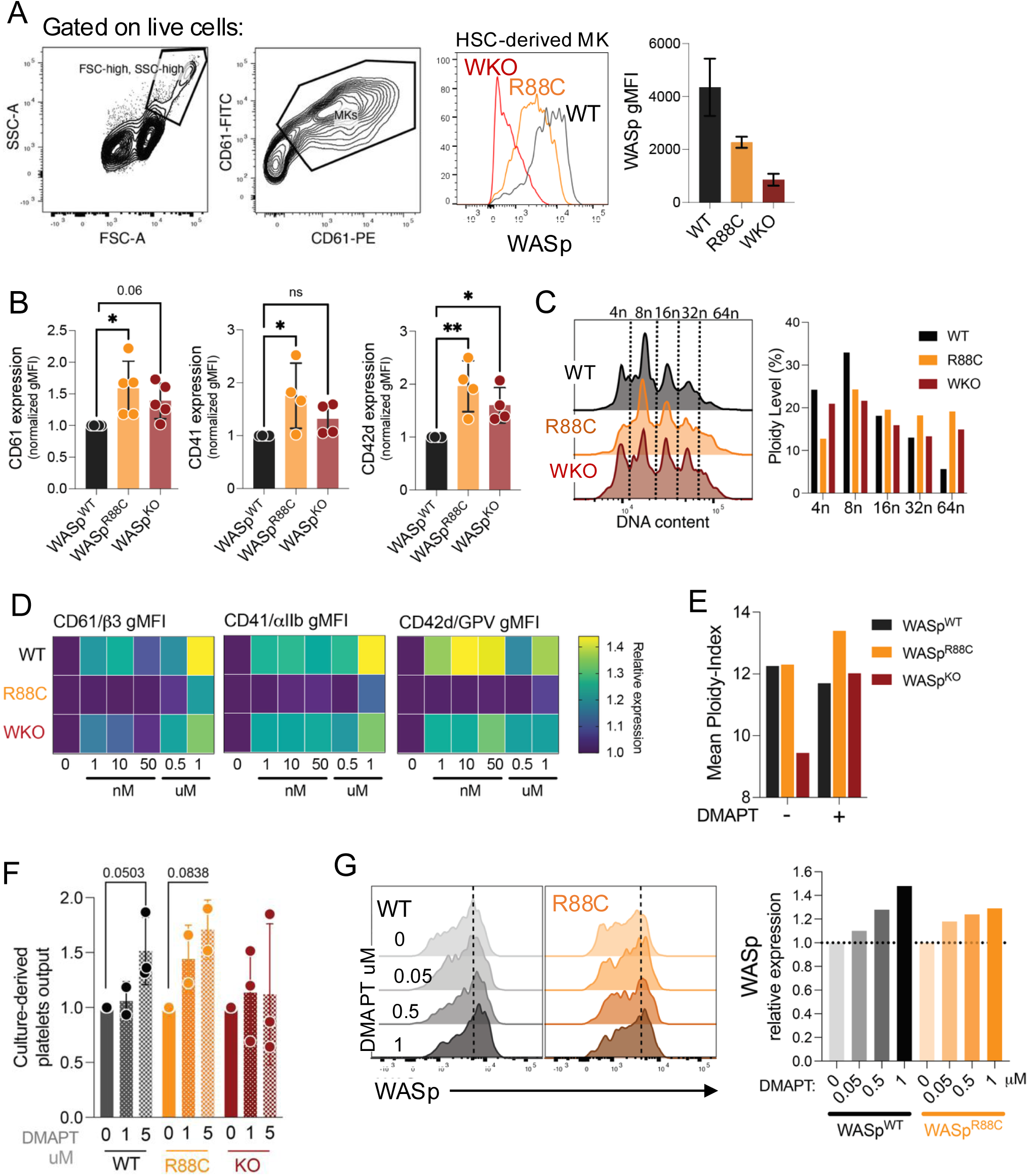
WAS murine primary megakaryocytes show increased maturation upon DMAPT treatment. **(A)** Representative gating strategy for MK and detection of intracellular WASp expression in primary HSC-expanded megakaryocytes derived from WT, WASp-R88C, and WASp-KO mice. Graph quantification shows WASp expression in MKs from the different genotypes. **(B)** Quantification of megakaryocyte maturation marker expression in WT, WASp-R88C, and WASp-KO primary megakaryocytes. Surface expression of CD61, CD41, and CD42d was assessed by flow cytometry and normalized to WT MKs cultured under same conditions**. (C)** Representative DNA-content histograms and quantification of ploidy distribution in WT, WASp-R88C, and WASp-KO primary megakaryocytes. **(D)** Representative histograms and quantification of megakaryocyte maturation markers following DMAPT treatment. **(E)** Quantification of mean ploidy index following DMAPT treatment. **(F)** Quantification of platelet output following DMAPT treatment. MKs were seeded in a fibrinogen-coated plate and incubated under agitation (160 RPM) for 24h. **(G)** Representative histograms and quantification of WASp expression in WT and WASp-R88C primary megakaryocytes treated with increasing concentrations of DMAPT. WASp expression was assessed by intracellular flow cytometry and normalized to WASp-KO. Statistical analysis: one-way ANOVA followed by Holm-Šídák’s multiple comparisons test. Experiments have at least n = 3 experimental replicates. Each dot represents one biological experiment.

When compared to WT cells, WAS-R88C and WASp-KO megakaryocytes had increased expression of CD61, CD41, and CD42d together with changes in ploidy distribution, suggesting dysregulated megakaryocyte maturation (Figure 4B–C; Supplementary Figure 6A, B). This altered maturation phenotype has been suggested in other studies (5,34).

Parthenolide has limited pharmacological properties, and more stable parthenolide analogs have been developed for translational studies (35,36). Subsequent primary megakaryocyte and *in vivo* studies were performed using DMAPT (dimethylamino-parthenolide), a parthenolide derivative with increased solubility and bioavailability (27,35). DMAPT treatment of primary WT, WAS-R88C, and WASp-KO murine megakaryocytes elevated expression of maturation markers CD61, CD41, and CD42d, increased megakaryocyte ploidy of WAS-R88C and WASp-KO cells, and stimulated platelets release in culture supernatants (Figure 4D-F, Supplementary Figure 6C, D). In addition, DMAPT increased intracellular WASp levels in WT and WAS-R88C primary megakaryocytes (Figure 4G; Supplementary Figure 6E). These findings show that parthenolide and its derivative DMAPT promote a coordinated megakaryocyte maturation program and enhance thrombopoietic output in MEG-01 cells and primary megakaryocytes. The effect of parthenolide treatment is detected in WT, class I WAS, and to a lesser extent, in class II WAS megakaryocytes.

### DMAPT boosts platelet counts *in vivo* in a class I WAS murine model

Based on that parthenolide treatment increased megakaryocyte maturation and *in vitro* platelet production, we set up an *in vivo* treatment protocol to examine whether DMAPT could enhance platelet production in murine WAS models. At baseline, WAS-R88C and WASp-KO mice had reduced platelet counts compared with WT mice, consistent with thrombocytopenia in WAS patients, whereas platelet size and complexity were similar to WT mice (Figure 5A, B). Mice were treated with DMAPT at 25 mg/kg, and platelet numbers were monitored at day 4, 7, and 10 (Figure 5C). DMAPT treatment increased circulating platelet counts in WT and WASp-R88C mice, reaching approximately a two-fold increase relative to baseline (Figure 5D-E, G). In contrast, WASp-KO mice did not show an enhanced platelet response after DMAPT treatment (Figure 5F, G). These results demonstrate that DMAPT enhanced platelet production *in vivo,* with a pronounced response in the WASp-R88C class I model. The absence of a clear platelet response in WASp-KO mice suggests that residual WASp expression contributes to the platelet-boosting effect of DMAPT.

**Figure 5:**
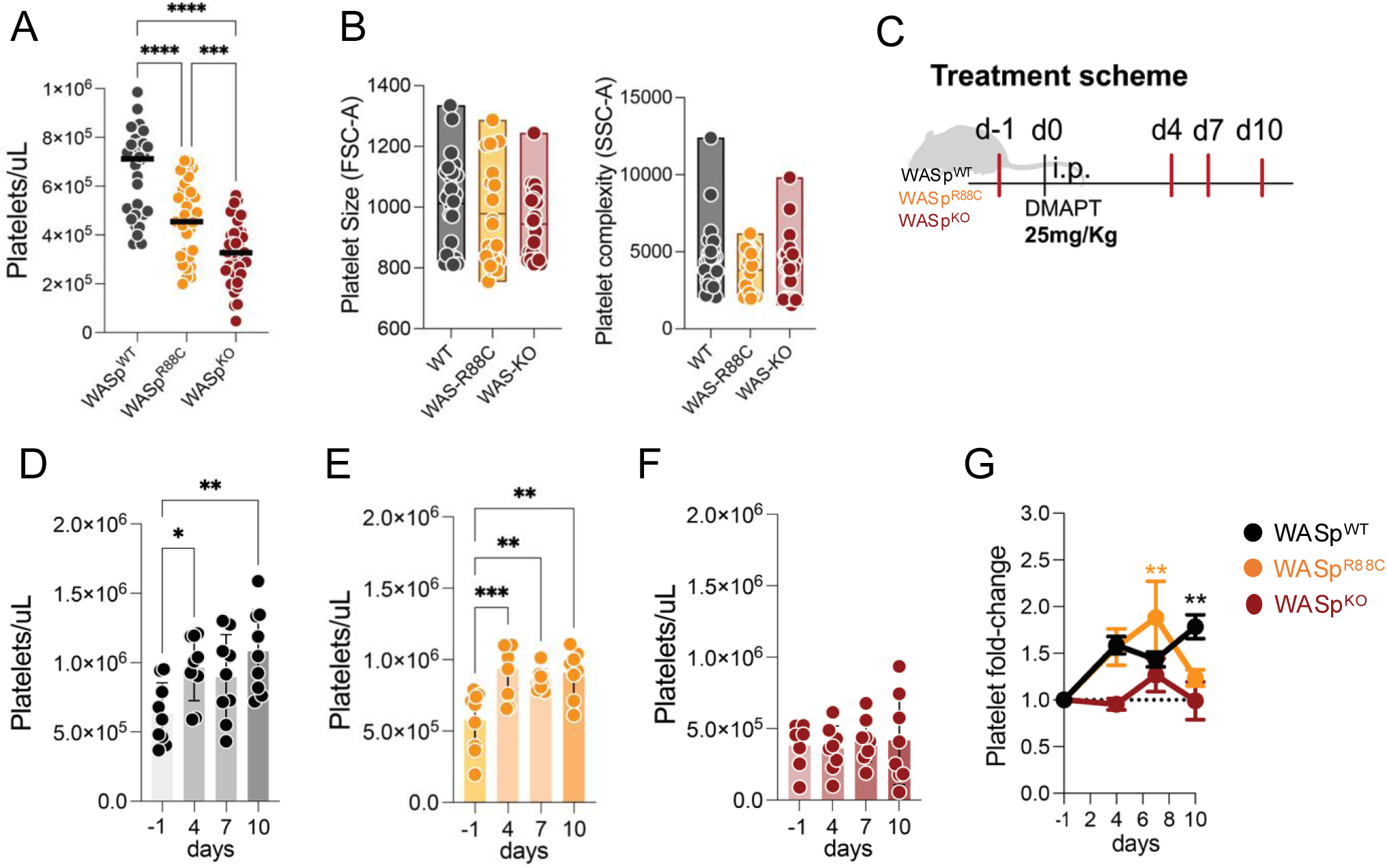
DMAPT increases platelet counts *in vivo* in WASp-R88C mice. **(A)** Scheme showing the strategy for generation of WASp-R88C mice. **(B)** Baseline peripheral blood platelet counts in whole blood of WT (n= 33), WASp-R88C (n= 33), and WASp-KO mice (n= 39). **(C)** Quantification of platelet size, given by geoMFI of FSC-A parameter (left) and platelet complexity given by geoMFI of SSC-A parameter (right). **(D)** Scheme showing treatment strategy. **(E-G)** Peripheral blood platelet counts in WT **(E)**, WASp-R88C **(F)**, and WASp-KO **(G)** mice following DMAPT treatment. Mice were treated with DMAPT at 25 mg/kg i.p., and platelet numbers were monitored at the indicated time points. **(H)** Fold-change in platelet counts relative to baseline for each genotype following DMAPT treatment. Fold-change was calculated for each mouse relative to its own day 0 platelet count. Each dot represents an individual mouse pooled from two independent experiments (D-F). Bars show mean ± SD. Statistical analysis: one-way ANOVA followed by Šídák’s (A). For longitudinal analyses (B–E), repeated-measures mixed-effects analysis was used with Dunnett’s multiple comparisons test comparing each time point with day 0 within each genotype.

### DMAPT enhances WASp and actin polymerization in activated human platelets and improves cytoskeletal defects in WAS patient platelets

To determine whether DMAPT directly affects actin regulation in human platelets, washed platelets from healthy donors were pre-incubated with DMAPT and stimulated with thrombin. Analysis showed that thrombin stimulation increased WASp, and this effect was further enhanced in DMAPT-treated platelets (Figure 6A, B). DMAPT-treated platelets displayed increased F-actin content following thrombin stimulation, compared with untreated controls (Figure 6A, B). These findings indicate that DMAPT promotes WASp accumulation and actin polymerization during platelet activation.

**Figure 6:**
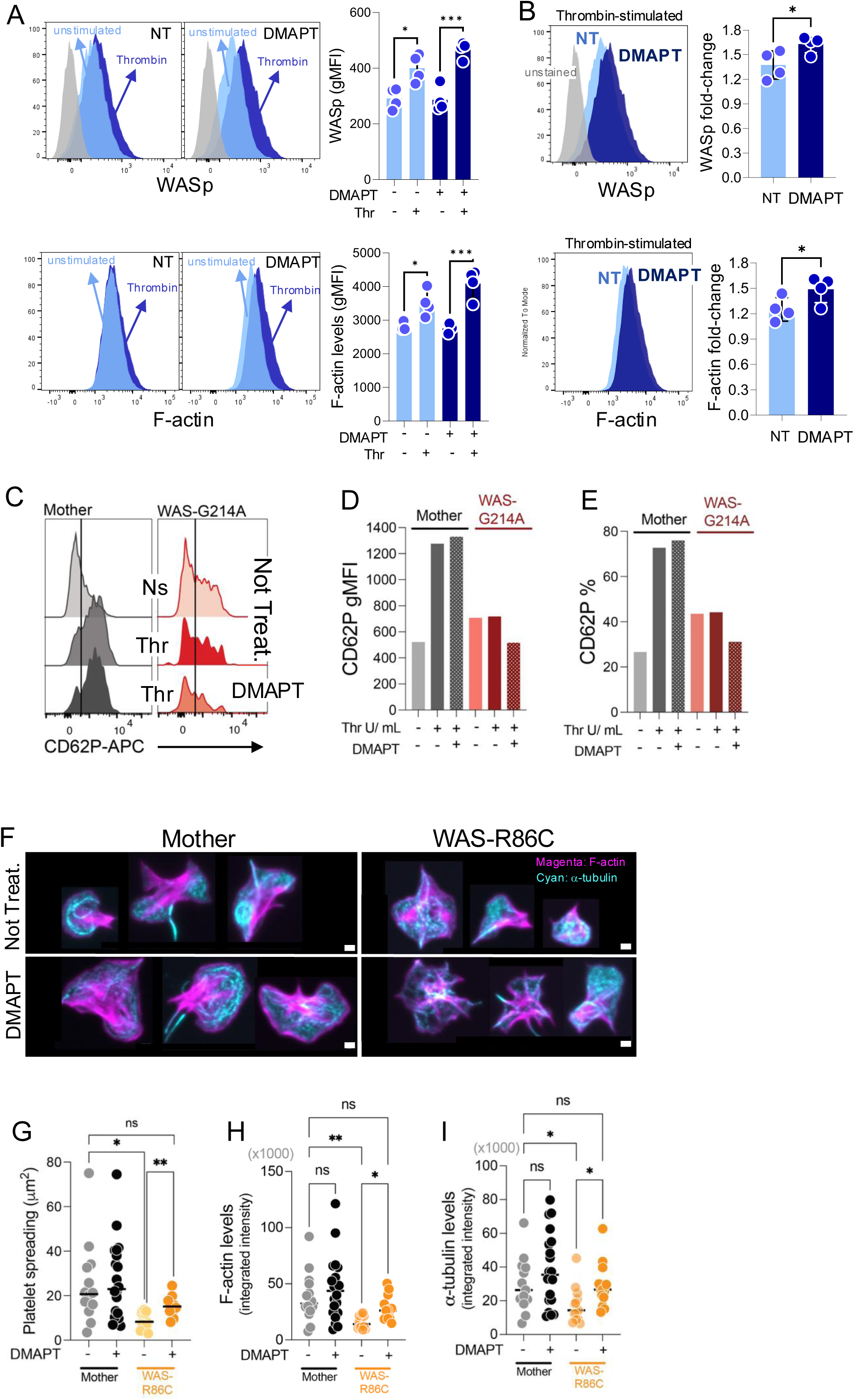
DMAPT enhances cytoskeletal responses in healthy donor platelets and partially improves cytoskeletal defects in WAS patient-derived platelets. **(A)** Representative flow cytometry histograms and graph quantification showing WASp expression (upper panel) and F-actin levels (bottom panel) in washed platelets from healthy donors following thrombin stimulation in the presence or absence of DMAPT. Unstimulated platelets are shown in light blue and thrombin-stimulated platelets in dark blue. **(B)** Quantification of WASp expression (upper penal) and F-actin levels (bottom panel) measured by flow cytometry. Data are presented as fold-change relative to untreated controls. Each symbol represents an independent donor. **(C)** Representative CD62P histograms of platelets from a WAS patient (red) carrying the G214A variant and the corresponding mother control (gray). Platelets were left untreated or incubated with DMAPT prior to thrombin stimulation. **(D-E)** Quantification of CD62P expression as gMFI (D) and frequency of CD62P-positive platelets (E) under the indicated conditions. **(F)** Representative confocal microscopy images of fibrinogen-spreaded platelets from a WAS patient carrying the R86C variant and the corresponding mother control treated with or without DMAPT. Platelets were allowed to spread for 10 minutes. F-actin (magenta) and α-tubulin (cyan) are shown. Scale bars, 1 μm. **(G-I)** Quantification of platelet spreading area (G), F-actin integrated fluorescence intensity (H), and α-tubulin integrated fluorescence intensity (I). Each dot represents independent donors (A, B) or a single platelet (G-I). Statistical analysis: one-way ANOVA followed by Šídák’s multiple comparisons test (A); Dunns multiple comparisons test (G-I); or paired t-test (B).

To investigate whether these effects could also be observed in WAS patient-derived platelets, we analyzed washed platelets from a patient carrying the WASp-G214A variant, a class II mutation leading to very low/absent WASp expression and microthrombocytopenia (Supplementary Figure 7). Platelets from the patient and the mother, used as control, were treated with DMAPT prior to thrombin stimulation, and platelet activation was assessed by CD62P surface expression. Consistent with a pre-activated phenotype, WASp-G214A platelets displayed elevated baseline CD62P expression compared with platelets from the maternal control (Figure 6C–E). Thrombin stimulation induced increased CD62P on platelets from the mother, but no difference to baseline was observed in the WASp-G214A platelets. DMAPT treatment had no further effect on CD62P on the maternal platelets but resulted in a modest reduction in both CD62P intensity and the frequency of CD62P-positive platelets from the WASp-G214A patient (Figure 6D, E). These data suggest that DMAPT may partially attenuate the increased basal activation status of WASp-G214A platelets. We next examined platelets from a patient carrying the class I WASp-R86C variant, associated with reduced WASp expression and microthrombocytopenia (Supplementary Figure 8A-D). Whereas the WASp-G214A patient platelets had only residual WASp expression, WASp-R86C platelets had WASp expression albeit lower than the maternal platelets. Moreover, basal CD62P expression was comparable between patient and maternal control platelets (Supplementary Figure 8E). Of note, similarly to the activation profile observed for WASp-G214A platelets, WASp-R86C platelets had a reduced activation response following thrombin stimulation, as evidenced by lower CD62P and PAC1 upregulation compared with control platelets (Supplementary Figure 8E, F). To evaluate the effects of DMAPT on platelet actin organization, washed platelets from the WASp-R86C patient and maternal control were allowed to spread on fibrinogen-coated coverslips in the presence or absence of DMAPT. Using confocal microscopy to evaluate platelet spreading and cytoskeletal remodeling, maternal platelets had increased spreading, higher F-actin and α-tubulin content when compared to WAS-R86C platelets. DMAPT treatment induced only minor changes of maternal platelets, whereas DMAPT-treated WAS-R86C platelets showed increased platelet spreading, and higher F-actin and α-tubulin staining intensity reaching the response of the maternal platelets (Figure 6F, G, H, I). Together, these results indicate that DMAPT partially improves cytoskeletal organization in WASp-R86C platelets. Notably, these observations are consistent with previous reports implicating parthenolide in the regulation of microtubule dynamics and tubulin modifications (37,38).

Finally, given the negative effects of parthenolide on cellular proliferation and survival of cancerous cells (39,40), we assessed whether DMAPT negatively influenced T cell responses of WAS patients. CD4⁺ T cells from healthy donors and patients (41) were stimulated through the T-cell receptor and cultured in the presence or absence of DMAPT. DMAPT treatment did not alter proliferation or expression of activation markers CD69 and CD25 in either healthy donor or patient-derived T cells (Supplementary Figure 9), indicating that DMAPT does not impair T cell activation under the conditions tested. Collectively, these findings indicate that DMAPT enhances cytoskeletal organization in human and WAS patient-derived platelets while not adversely affecting T cell activation or proliferation under the conditions tested.

## Discussion

Microthrombocytopenia of WAS patients remain a therapeutic challenge, particularly for individuals who carry class I WAS gene variants and do not qualify for gene therapy or hematopoietic stem cell transplantation (20). Here, we established a flow cytometry-based drug screening to identify small molecules capable of increasing WASp abundance in megakaryocytic cells expressing degradation-prone WAS variants. This approach identified parthenolide as a candidate compound that promotes megakaryocyte maturation and enhances platelet production in WAS models. Using both *in vitro* and *in vivo* systems, including murine models and patient-derived samples, we show that parthenolide-based treatment improved thrombopoietic responses and may represent a therapeutic strategy for WAS-associated thrombocytopenia.

The mechanisms underlying WAS-associated thrombocytopenia are complex and likely involve both peripheral platelet clearance and intrinsic defects in platelet production. Several studies support the idea that the platelet defect in WAS begins, at least in part, at the megakaryocyte level. WASp-deficient megakaryocytes have abnormal responses to bone marrow extracellular matrix cues, impaired migration, loss of actin-rich podosomes, and premature proplatelet or platelet release within the bone marrow compartment rather than at the vascular niche (5,42,43). Such ectopic platelet production could reduce efficient platelet entry into circulation and contribute to thrombocytopenia. In addition, WAS thrombocytopenia has been linked to peripheral platelet defect and immune dysregulation, including autoreactive B-cell responses and antiplatelet antibodies (6,8,44,45). Together, these observations highlight the need for therapies that improve both megakaryocyte output and platelet function. Our data support parthenolide/DMAPT as an interesting thrombopoietic candidate. In megakaryocytes, parthenolide/DMAPT enhanced ERK1/2 phosphorylation, upregulated CD41, CD61, and CD42d expression, increased ploidy, and promoted adhesion-induced spreading on fibrinogen, all indicators of final maturation of megakaryocytes and functional priming for proplatelet formation (28,34,35). Importantly, these maturation-associated changes were accompanied by increased platelet output *in vitro*. For human platelets, parthenolide/DMAPT pretreatment increased detectable intracellular WASp and amplified F-actin polymerization upon activation. In patient-derived samples, parthenolide/DMAPT pretreatment improved platelet spreading, suggesting a rapid effect on actin-dependent platelet responses.

The mechanism by which parthenolide/DMAPT promotes megakaryocyte maturation is likely to be multifactorial. Although the screening strategy was initially designed to identify compounds that elevated WASp abundance in cells expressing unstable class I variants, the functional effects observed in WASp-KO megakaryocyte models suggest that parthenolide/DMAPT activity cannot be explained solely by direct WASp expression. This is consistent with the known pleiotropic biology of the drug (27,37,38,46–54). Parthenolide directly binds and inhibits IκB kinase, thereby suppressing NF-κB activation (27,46,54). In fact, a previous study show that parthenolide increased platelet-like particle production from MEG-01 and MO7e cells and enhanced platelet production from primary mouse and human megakaryocytes, with evidence that this effect was associated with NF-κB inhibition (27). Parthenolide inhibits IL-6-family cytokine signaling by covalently targeting JAK family kinases and by inhibition of STAT3 Tyr705 phosphorylation (49)(51). These pathways are relevant because inflammatory and cytokine signaling networks can influence megakaryocyte maturation and platelet production.

In addition to effects on inflammatory signaling, parthenolide may influence cytoskeletal pathways relevant to megakaryocyte biology. Megakaryocyte maturation and platelet formation require extensive actin and microtubule remodeling, including cytoplasmic expansion, demarcation membrane system organization, proplatelet elongation, and platelet release (31,32). Parthenolide/DMAPT interferes with tubulin and microtubule organization, including stimulation of tubulin assembly *in vitro* and alteration of microtubule architecture in cells (this study)(37). In neuronal cells, parthenolide/DMAPT inhibits microtubule detyrosination and promotes axon growth, highlighting a biologically active connection between this compound class and microtubule dynamics (38). We show here that parthenolide improved the actin cytoskeleton remodeling and induced increased spreading of WAS platelets. This shows a broader effect of parthenolide on the cell cytoskeleton where it can affect microtubule and actin polymerization dynamics, both critical for megakaryocyte production of platelets and platelet function (31,32).

One important observation from our *in vivo* studies is that DMAPT elevated platelet counts in WT and WASp-R88C mice but did not produce a comparable platelet increase in WASp-KO mice. This result may appear inconsistent with the *in vitro* observations showing pro-maturation effects of WASp-KO megakaryocytes. This may reflect the difference between cell-intrinsic maturation readouts in cell cultures and productive platelet release into the circulation *in vivo*. In the bone marrow, megakaryocytes must coordinate maturation, migration, positioning at sinusoidal vessels, proplatelet extension into the vascular space, and platelet release under the influence of extracellular matrix, stromal cells, sinusoidal endothelium, chemokines, and shear forces (55,56). WASp-deficient megakaryocytes have premature platelet release within the marrow compartment (5). Therefore, parthenolide/DMAPT may be sufficient to enhance maturation-associated processes *in vitro* but insufficient to overcome the combined *in vivo* requirements for megakaryocyte positioning, cytoskeletal polarity, vascular platelet release, and platelet survival in complete WASp deficiency. This interpretation is also consistent with the idea that class I WAS variants may preserve enough residual WASp to allow pharmacological boosting of thrombopoiesis, whereas WASp-KO megakaryocytes may lack essential cytoskeletal functions required for efficient platelet release into the circulation. Thus, class I WAS variants may be particularly responsive to interventions that combine increased maturation signaling with partial restoration or preservation of cytoskeletal function.

The broader implication of our findings may extend beyond WAS. Several inherited thrombocytopenias arise from defects in actin cytoskeleton regulation, highlighting the importance of cytoskeletal remodeling for megakaryocyte maturation, proplatelet formation, and platelet function. Deletion of CDC42-interacting protein 4 (CIP4), a WASp-interacting regulator of membrane and actin remodeling, causes thrombocytopenia and impaired proplatelet formation (57). ARPC1B deficiency, which disrupts the Arp2/3 complex downstream of WASp-dependent actin nucleation, is associated with microthrombocytopenia and defective proplatelet formation (58). These observations suggest that parthenolide/DMAPT may be relevant not only for WAS, but also for a broader group of immune actinopathies and inherited thrombocytopenias characterized by impaired cytoskeletal remodeling (59). Current pharmacological approaches to increasing platelet counts largely rely on stimulation of the thrombopoietin-MPL axis by TPO receptor agonists such as romiplostim and eltrombopag. This treatment increases platelet counts in some patients with inherited thrombocytopenia, including WAS, but responses are variable and functional defects may not be corrected (21–23,60,61). Therefore, compounds that act downstream or parallel to c-MPL, rapidly promoting maturation or release programs, may complement existing therapies or provide alternative approaches for thrombocytopenia where TPO receptor signaling is insufficient or where platelet production is limited. A potential translational advantage of parthenolide-based treatment is the apparent ability to influence platelet output over a relatively short time frame. This possibility could be clinically relevant in settings requiring more rapid improvement in platelet production, such as perioperative management, bleeding risk reduction, or bridging to definitive therapy.

### Limitations and Future Directions

Parthenolide contains a reactive functional group capable of targeting cysteine residues (46,50,62) and its precise mechanism of action in megakaryocytes and its protein target(s) remain to be fully defined. It will be valuable to explore the structure–activity relationship around the core parthenolide scaffold to optimize specificity and therapeutic potential. We detected increased platelets upon DMAPT treatment *in vivo* in WAS-R88C mice which is encouraging. Several more steps would be necessary to complete before translating treatment into WAS patients with thrombocytopenia. While the anti-inflammatory action of parthenolide/DMAPT likely would be beneficial in WAS (6), this property may have different consequences in other thrombocytopenic conditions. Because parthenolide modulate platelet activation, careful dose optimization will be required to enhance platelet output without impairing platelet hemostatic function.

## Acknowledgement

We thank the Chemical Biology Consortium Sweden (CBCS) at SciLifeLab and Karolinska Institutet for access to compound libraries. CBCS is a national research infrastructure funded by the Swedish Research Council (dr.nr.2021-00179) and Science for Life Laboratory in Sweden (www.scilifelab.se). Special thanks to Anna Eriksson of CBCS Umeå University for her guidance in assay development and insightful discussions. We thank the Karolinska Genome Engineering and the Karolinska transgenic animal core facility for generation of the WAS-R88C mice. We are grateful to Israeli WAS Association (IWASA, Israel) and the Wiskott–Aldrich Syndrome foundation (WAF, USA) for their support and advice.

## Authorship Contributions

R.C.V. and L.S.W. was responsible for study conceptualization and planning, and supervision of students and wrote the original draft of the manuscript; R.C.V. was responsible for cell line model generation, flow cytometry-screening assay development, experimental procedures, data acquisition, and data analysis and interpretation; L.G.P. was responsible for cell lines model generation, and technical support; T.Y., M.H., G.Z., M.D.G., E.J.V. and I.R.E. were responsible for experimental procedures; A.L.G. was responsible for technical advice and for providing access to the CBCS drug screening libraries; O.E. was responsible for patient contact and shipment of blood samples; L.S.W. was responsible for project supervision; and all authors reviewed and edited the manuscript.

This work was supported by project grants from the IWASA to R.C.V. and L.S.W., a postdoctoral fellowship from Wenner-Gren Foundations to L.G.P., a KID KI PhD fellowship to T.Y., the Swedish Research Council, Cancer Society, Worldwide Cancer Research, Childhood Cancer Fund, Radiumhemmet Research Funds, and Karolinska Institutet to L.S.W.

## Supplementary Figure legends

**Supplementary Figure 1: CRISPR/Cas9-mediated knockout of WASp. (A)** Flow cytometry histograms showing WASp protein expression in individual clones generated using gRNA1 (targeting exon 7, left) or gRNA2 (targeting exon 2, right). Scramble control and secondary antibody-only (SecAb) controls are shown for comparison. Percentages indicate the proportion of WASp-positive cells. Right panels show geometric mean fluorescence intensity (geoMFI) ratios of WASp expression for each clone, normalized to the secondary antibody background. Bars in red indicate clones with nearly complete WASp loss. **(B)** Schematic representation and sequencing chromatogram of a detected insertion mutation in a clone edited with gRNA2, introducing a duplication of Lys81_Ser82 (p.Lys81_Ser82insLys), leading to reduced WASp expression. **(C)** Schematic of the WASp gene segments and corresponding point mutations. Left: R86C mutation (c.290C>T) changes the first nucleotide of the codon for arginine (Arg) to encode cysteine (Cys). Right: L39P mutation (c.150T>C) alters the second nucleotide of the codon for leucine (Leu) to encode proline (Pro). Top rows show the original wild-type sequences and translations; middle rows display the aligned amplicon sequences with the mutation site highlighted; bottom rows show representative Sanger sequencing chromatograms confirming the mutations.

**Supplementary Figure 2: MEG-01 EGFP-expressing WASp mutants. (A)** Flow cytometry analysis of WASp expression and EGFP reporter signal in cells expressing WT, L39P, or R86C variants. Histograms (left) show WASp and EGFP signal distributions. Bar graphs (right) represent gMFI values of WASp and EGFP for each variant. Data is shown as mean ± SD from independent experiments. **(B)** MEG-01 cells expressing WT, L39P, or R86C WASp-EGFP fusion proteins were treated with cycloheximide (CHX, 100 μM) to inhibit new protein synthesis, and EGFP fluorescence was monitored over 24 hours. Left: Quantification of EGFP gMFI, normalized to untreated (NT) controls for each construct. Right: Representative flow cytometry histograms showing EGFP signal decay over time for WT, L39P, and R86C variants.

**Supplementary Figure 3: Flow cytometry-based drug screening assay development and optimization using MEG-01 expressing the WASp-R86C class I variant.** For assay development the cells were tested in different conditions (treated or not with CHX) and assays assessed the best performance regarding Z-factor. **(A)** Scheme of 96-well plate setup (left) and heat maps indicating the number of cells acquired per well in non-treated (middle) or CHX-treated (right) incubated overnight. **(B)** Representative histograms showing EGFP signal in cells treated or not with proteosome inhibitors MG-132 or BTZ. Upper panel: cells culture without CHX; bottom panel: cells cultured with CHX. Vertical dotted line indicates mean signal values of negative control. **(C)** WASp accumulation-index (AI) quantification based on EGFP gMFI in cells cultured or not with CHX. **(D)** Z-factor values of the screening assay in different conditions. **(E)** Scheme of 96-well plate layout for the screening assay. The FACS-based screening was performed under CHX treatment. **(F)** Representative gating strategy and histograms showing EGFP signal intensity in negative, bortezomib-treated positive control (BTZ), and a representative low and high-AI sample. **(G)** Representative dot plot (left) and quantification graph (right) showing WASp-AF647 signal in cells treated or not with BTZ; all cells were under CHX *de novo* translation inhibition. **(H)** Accumulation-index (AI) of samples shown in A indicate the dual WASp detection by EGFP and WASp-AF647 signals; Z-factor values are indicated in the graph. **(I)** Quantification graphs of WASp staining in EGFP-expressing WASp WT, WASp-L39P, and WASp-R86C cells showing the reduced WASp degradation in cultures treated with #08 (parthenolide) or #25 (auranofin).

**Supplementary Figure 4: Screening of compounds from the SPECS library. (A)** EGFP accumulation-index (AI) scatter plot from primary screening of SPECS library. Points above threshold (gray zone) indicate 60 selected positive compounds. WASp-R86C cells used. **(B)** Counter-screening of the 60 hits in MEG-01 WASp-insK82 cells, measuring WASp accumulation-index after 48h incubation. Blue area indicates selection threshold (AI > 1.4); 9 compounds were positively selected. **(C)** WASp accumulation-index validation of the final 4 selected compounds (SU11274, BRD-7389, AMG-548, and Ro-106-9920) at indicated concentrations, comparing MEG-01 WASp-insK82 cells (orange background) to MEG-01 WKO cells (red background). Points represent independent replicates. **(D)** Western blot validation of WASp accumulation in WASp-R86C cells treated with indicated compounds under CHX conditions. GAPDH used as loading control. Bottom graph shows quantification of WASp relative expression normalized to GAPDH.

**Supplementary Figure 5: Compound toxicity and PMA-induced platelet production. (A)** Representative flow cytometry plots of DNA content (DAPI) versus CD41 expression for WT, insK82, and WKO MEG-01 cells treated or not (NT) with PMA (25 ng/mL, 48h). Percentages indicate cells within 4n and ≥8n ploidy gates in the CD41⁺ population. **(B)** Quantification of CD41⁺ platelet-like particles released by WT, insK82, and WKO cells culture supernatant in NT or PMA-treated condition. **(C)** DNA content analysis (DAPI staining) of CD41-negative populations from WT, insK82, and WKO MEG-01 cells treated or not (NT) with PTW08 (10µM, 48h). Left: Representative histograms. Right: Quantification graphs showing percentage of cells with 4n and ≥8n ploidy. **(D)** Representative FSC-A versus SSC-A flow cytometry contour plots illustrating morphology and granularity changes of WT, insK82, and WKO MEG-01 cells following PTW08 treatment compared to NT controls. Statistical analysis: One-way ANOVA followed by Šídák’s multiple comparisons test (B).

**Supplementary Figure 6: Hematopoietic stem cell-derived megakaryocytes analysis. (A)** Flow cytometric quantification of megakaryocyte differentiation markers CD61, CD41, and CD42d in megakaryocytes derived from WASp^WT^, WASp^R86C^, and WASp^KO^ cultures. Data are presented as geometric mean fluorescence intensity (gMFI). Each symbol represents an independent experiment. **(B)** Representative flow cytometry gating strategy used to identify mature megakaryocytes and assess ploidy. Viable cells were selected based on Live/Dead exclusion, followed by FSC-A and CD61 expression gating. Mature megakaryocytes were identified as CD61^+^CD42d^+^ cells, and DNA content was determined by DAPI staining to quantify ploidy levels (2N-64N). Representative histograms and contour plots are shown. **(C)** Representative ploidy histograms and quantification of ploidy distribution in WASp^WT^, WASp^R86C,^ and WASp^KO^ megakaryocyte cultures treated with vehicle (NT), 0.5 μM DMAPT, or 1 μM DMAPT. The frequency of cells within each ploidy class is shown on the right. **(D)** Quantification of culture-derived platelet production from WASp^WT^, WASp^R86C,^ and WASp^KO^ megakaryocyte cultures treated with 0, 1, or 5 μM DMAPT. Data are presented as the number of culture-derived platelets per mL. Each symbol represents an independent experiment. **(E)** WASp expression levels measured by flow cytometry in WASp^WT^, WASp^R86C,^ and WASp^KO^ megakaryocytes cultured in the presence of the indicated concentrations of DMAPT. Data are presented as geometric mean fluorescence intensity (gMFI).

**Supplementary Figure 7: Phenotypic characterization of platelets from a patient carrying the WASp-G214A variant. (A)** Representative flow cytometry gating strategy used for platelet identification in samples from the mother (control) and the WASp-G214A patient. Platelets were selected based on forward and side scatter properties and further identified by CD41 expression. The percentage of platelet events within the selected gates is indicated. **(B)** Representative contour plots showing platelet size and complexity distribution (FSC-A versus SSC-A) in mother control and WASp-G214A patient platelets. Histograms on the top and right illustrate FSC-A and SSC-A signal distribution, respectively, showing reduced platelet size (FSC-A) on patient. **(C)** Representative flow cytometry histograms showing intracellular WASp expression in platelets from the mother (grey) and the WASp-G214A patient (red). **(D)** Quantification of circulating platelet counts in the maternal control and WASp-G214A patient. Data are presented as platelets/μL blood.

**Supplementary Figure 8: Phenotypic characterization of platelets from a patient carrying the WASp-R86C variant. A)** Representative flow cytometry gating strategy used for platelet identification in samples from the mother (control) and the WASp-R86C patient. Platelets were selected based on forward and side scatter properties and further identified by CD41 expression. The percentage of platelet events within the selected gates is indicated. **(B)** Representative contour plots showing platelet size and complexity distribution (FSC-A versus SSC-A) in mother control and WASp-R86C patient platelets. Histograms on the top and right illustrate FSC-A and SSC-A signal distribution, respectively, showing reduced platelet size (FSC-A) on patient. **(C)** Representative flow cytometry histograms showing intracellular WASp expression in platelets from the mother (grey) and the WASp-R86C patient (red). **(D)** Quantification of circulating platelet counts in the maternal control and WASp-R86C patient. Data are presented as platelets/μL blood. **(E)** Representative flow cytometry histograms showing platelet activation following stimulation with increasing concentrations of thrombin (0, 0.1, and 0.5 U/mL). Activation was assessed by CD62P surface expression (upper panel) and PAC1 binding (bottom panel), a marker of activated integrin αIIbβ3, in maternal control (grey) and WASp-R86C (red) platelets. **(F)** Quantification of CD62P and PAC1 expression measured as geometric mean fluorescence intensity (gMFI) after 2 and 5 min of thrombin stimulation. Maternal control and WASp-R86C platelets were stimulated with the indicated thrombin concentrations (0, 0.1, and 1 U/mL). Data show a reduced activation response in WASp-R86C platelets compared with control platelets.

**Supplementary Figure 9: DMAPT does not impair activation or proliferation of CD4⁺ T cells from healthy donors and WAS patients. (A)** Representative flow cytometry gating strategy used for analysis of CD4⁺ T cells. Cells are part of Lab-Biobank, were previously sorted and expanded CD4⁺ T cells blast and cryopreserved. After thawing, cells were kept in IL-7 (20ng/mL) and IL-15 (2ng/mL), and harvest 3-days after stimulation. **(B)** Representative CD69 and CD25 expression in unstimulated and ImmunoCult™-stimulated CD4⁺ T cells. **(C)** Representative CellTrace Violet (CTV) dilution profiles showing proliferation of unstimulated and ImmunoCult™-stimulated CD4⁺ T cells. **(D)** Representative histograms of CD25 and CD69 expression following ImmunoCult™ stimulation in CD4⁺ T cells from the age-matched healthy control S3 (sister control of patient R268W), an unrelated healthy donor (HD), and WAS patients carrying the G214A and I294T variants. **(E-F)** Quantification of CD25 and CD69 expression measured as geometric mean fluorescence intensity (gMFI) in healthy donor and WAS patient-derived CD4⁺ T cells cultured in the presence of increasing concentrations of DMAPT, as indicated. DMAPT treatment did not substantially alter expression of either activation marker compared with stimulated untreated controls. **(G)** Representative CTV proliferation profiles of CD4⁺ T cells from S3, healthy donor (HD), and WAS patients stimulated with ImmunoCult™ and cultured in the presence of increasing concentrations of DMAPT (0, 1, 0.1, 0.01, and 0.001 μM). Comparable CTV dilution profiles were observed across all conditions, indicating that DMAPT does not impair T-cell proliferation compared to its own not-treated condition.

