## Supplementary Figures for "Parthenolide boosts megakaryocyte maturation and restores platelet responses in Wiskott–Aldrich syndrome"

Supplementary Figure 1

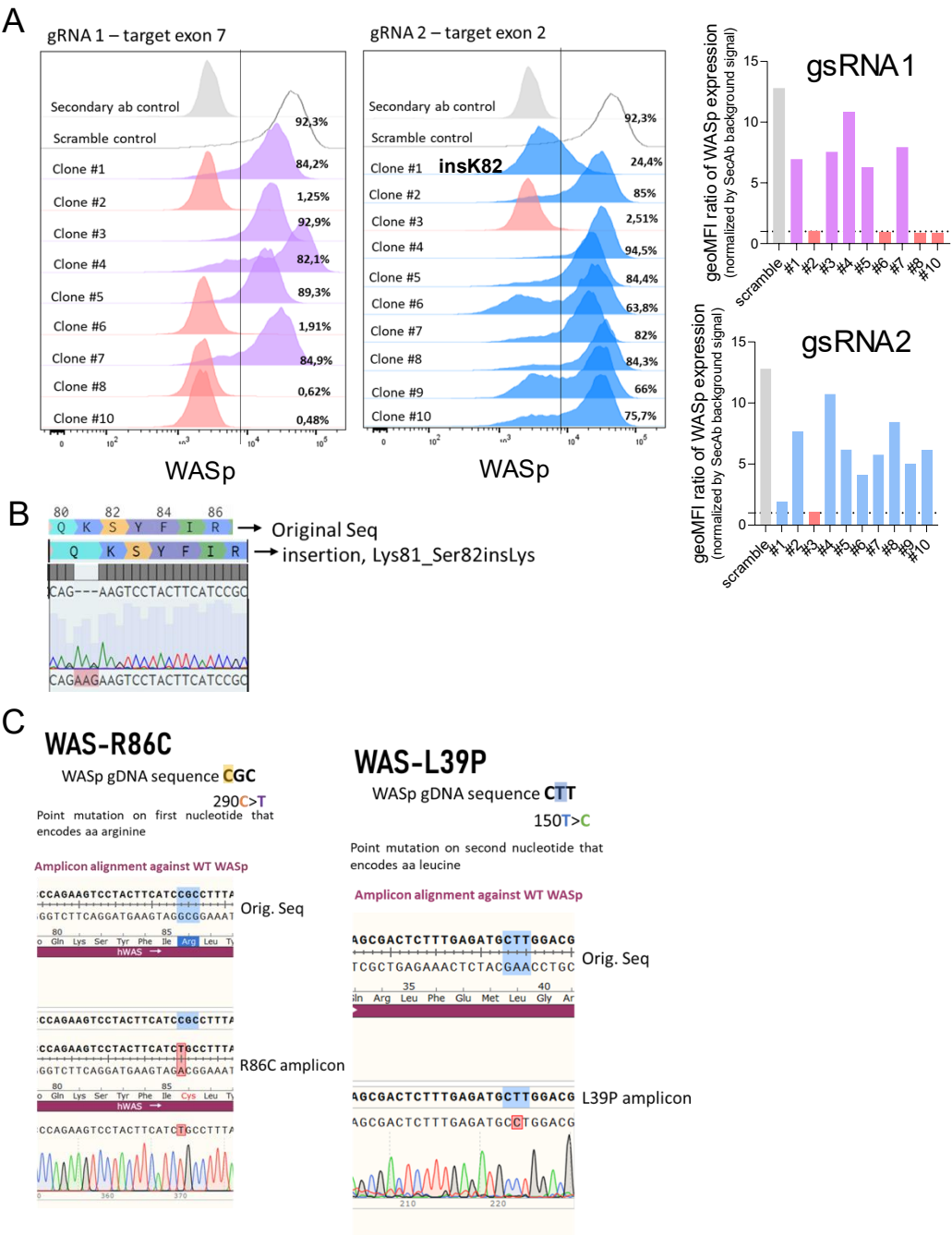

Supplementary Figure 2

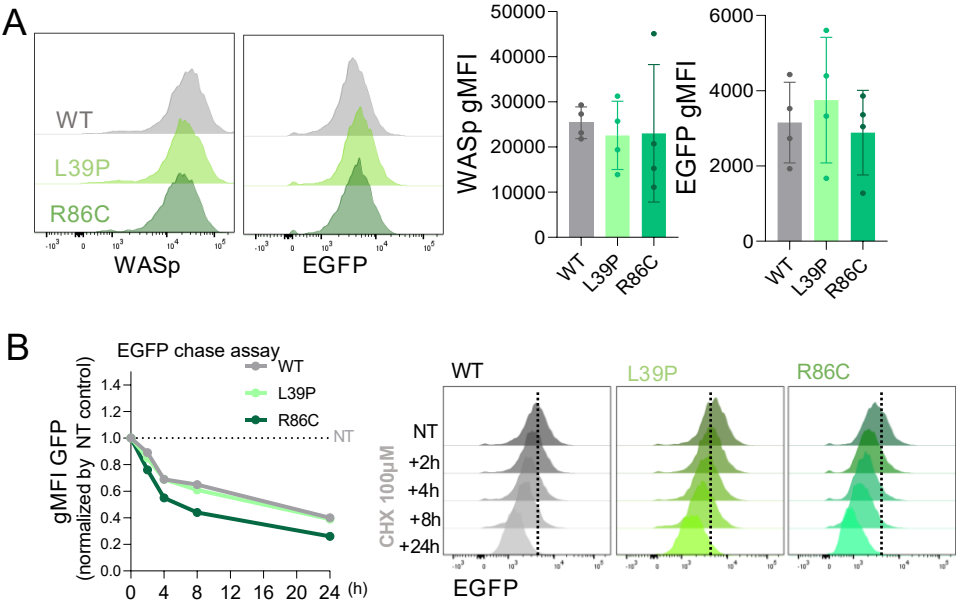

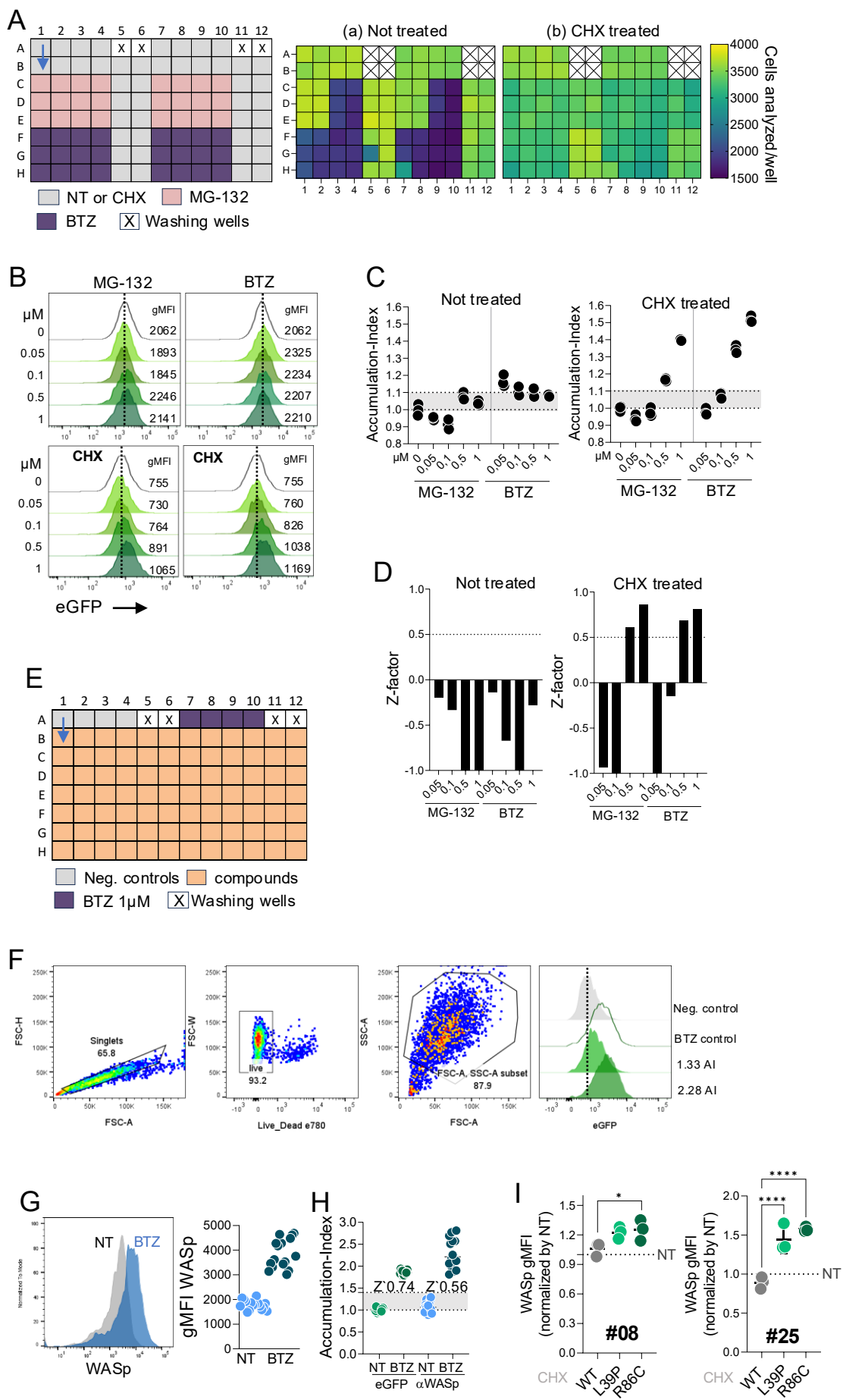

Supplementary Figure 4

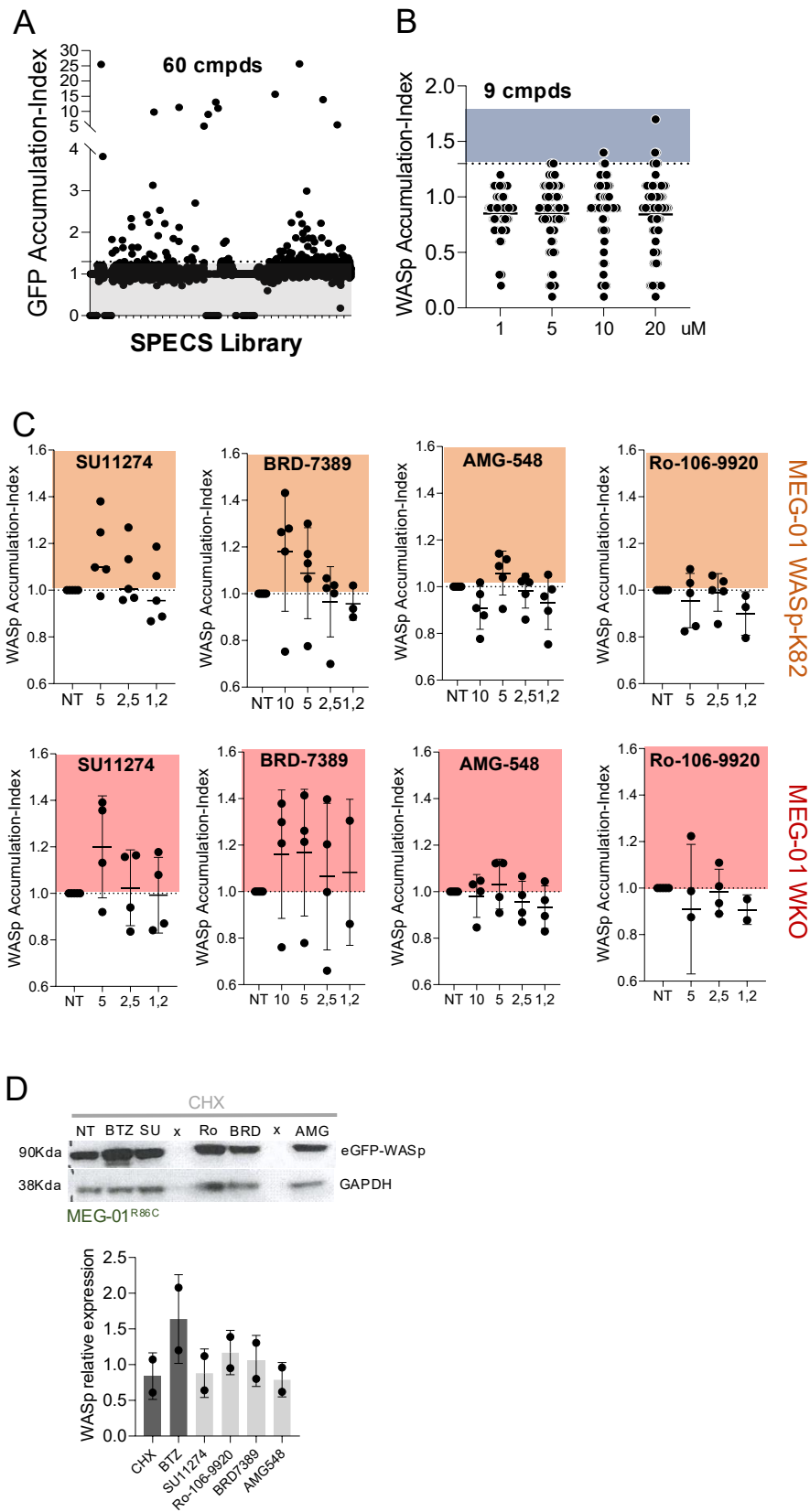

Supplementary Figure 5

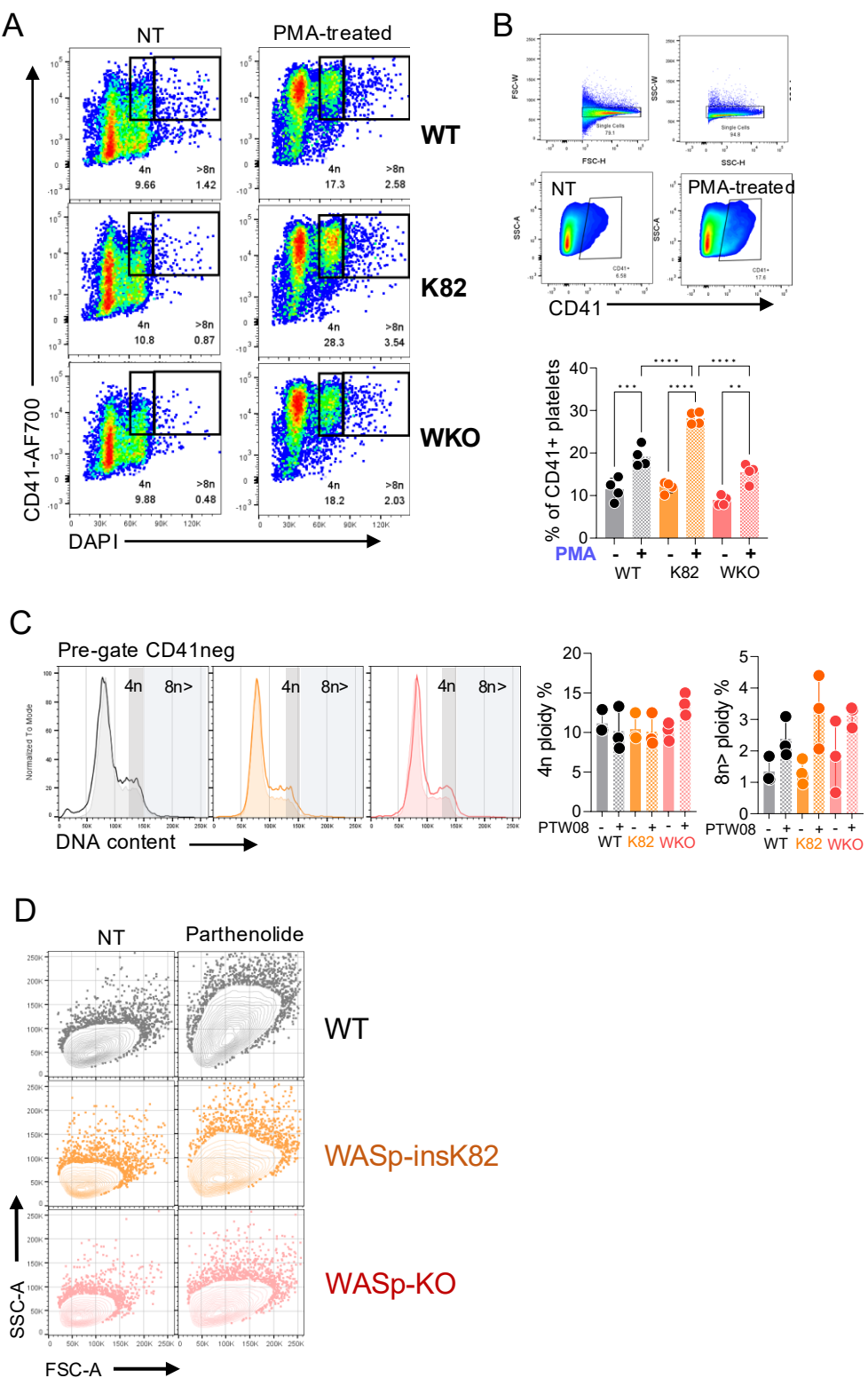

Supplementary Figure 6

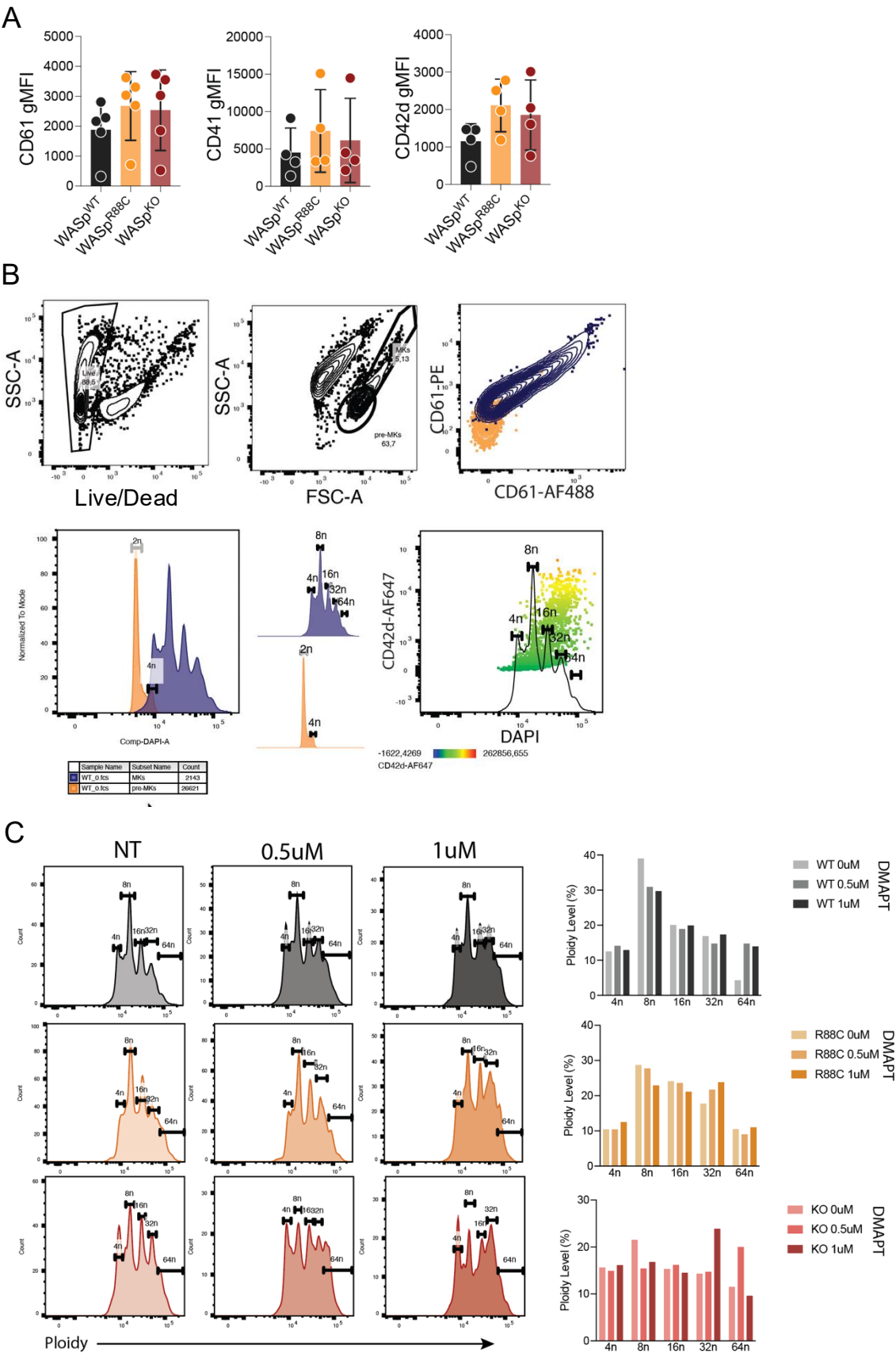

Supplementary Figure 6

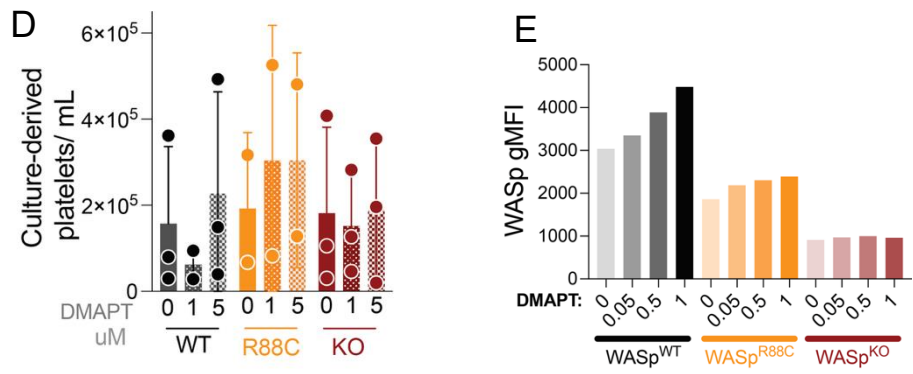

Supplementary Figure 7

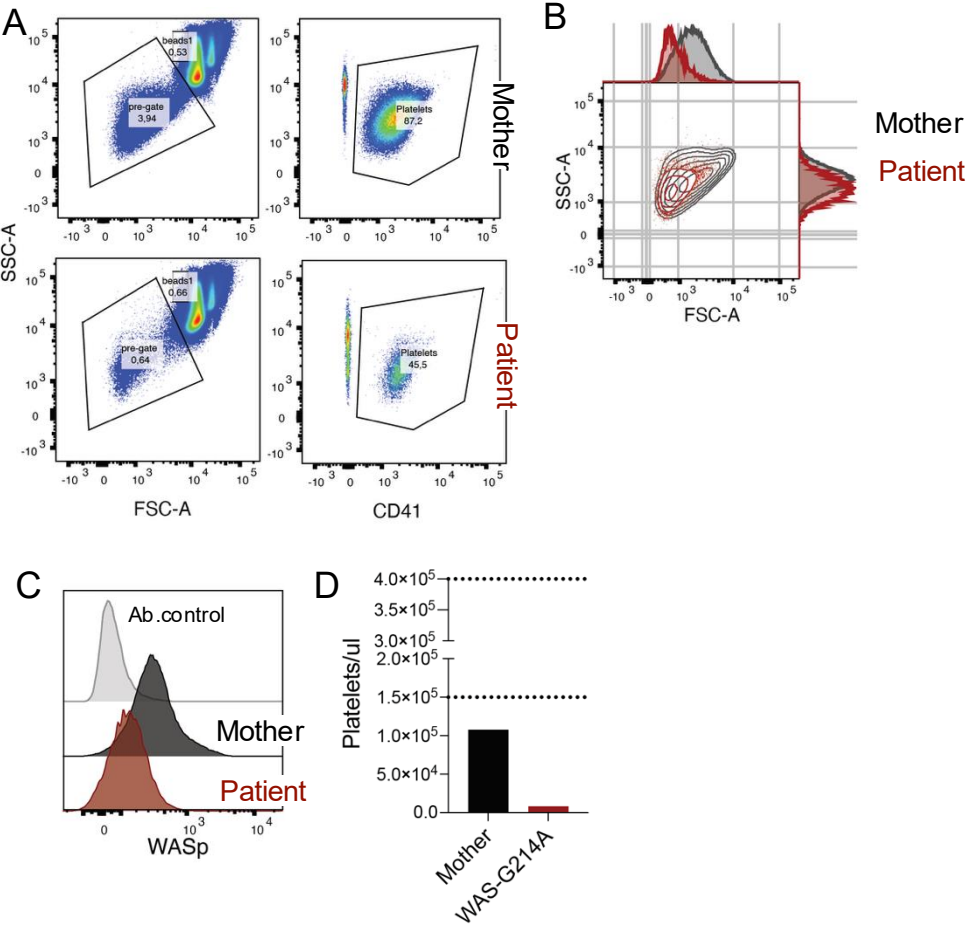

Supplementary Figure 8

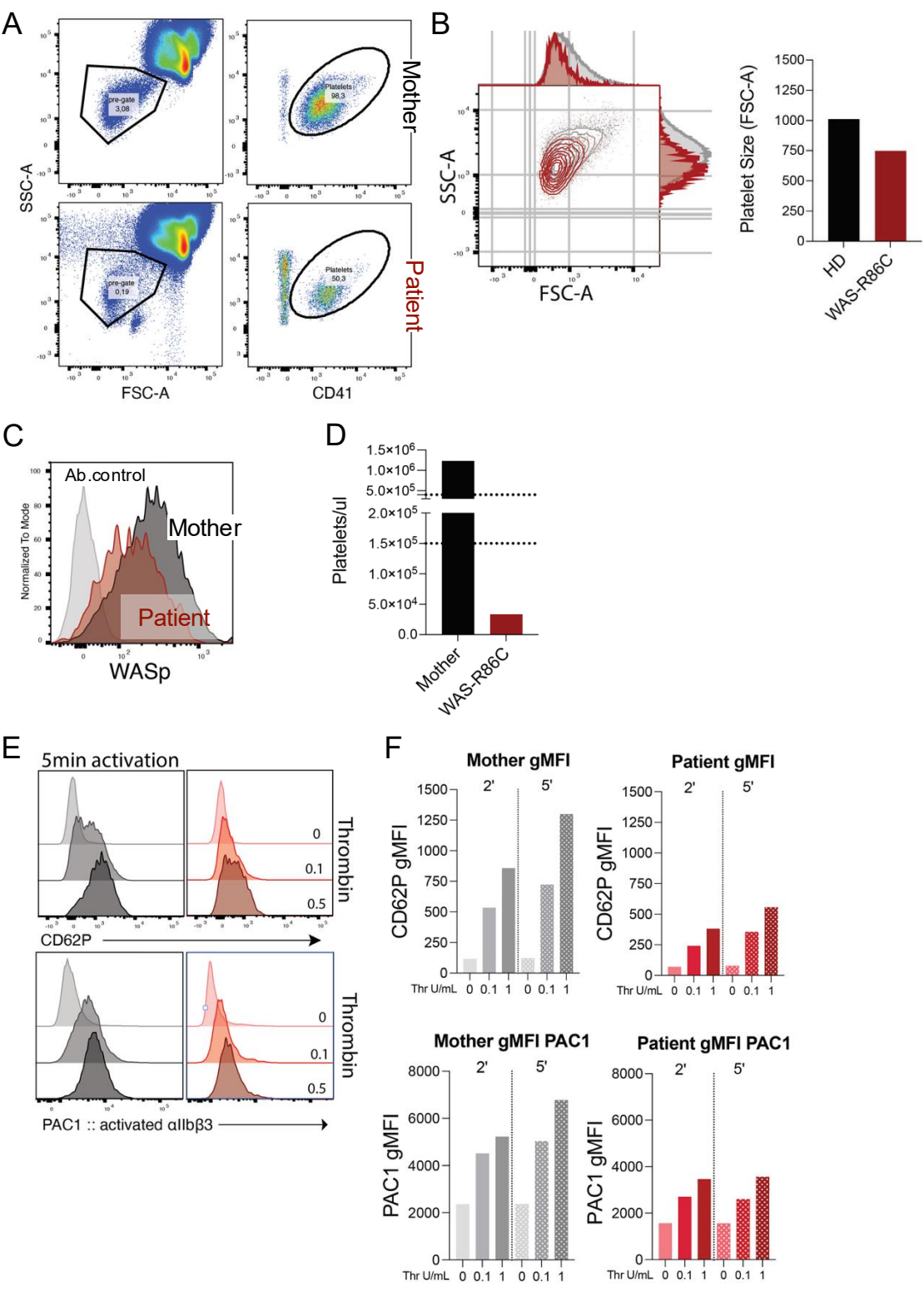

Supplementary Figure 9

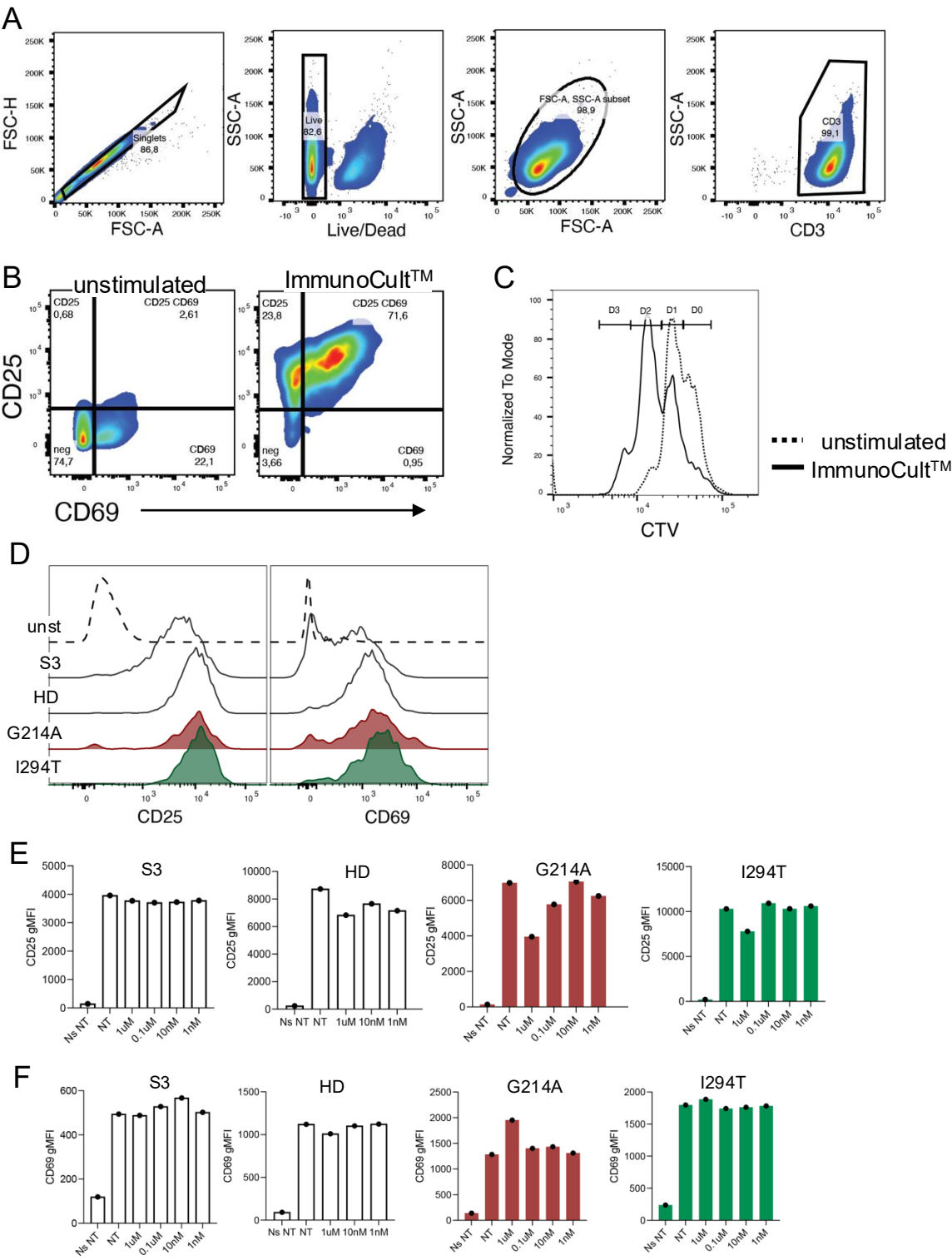

Supplementary Figure 9

## G

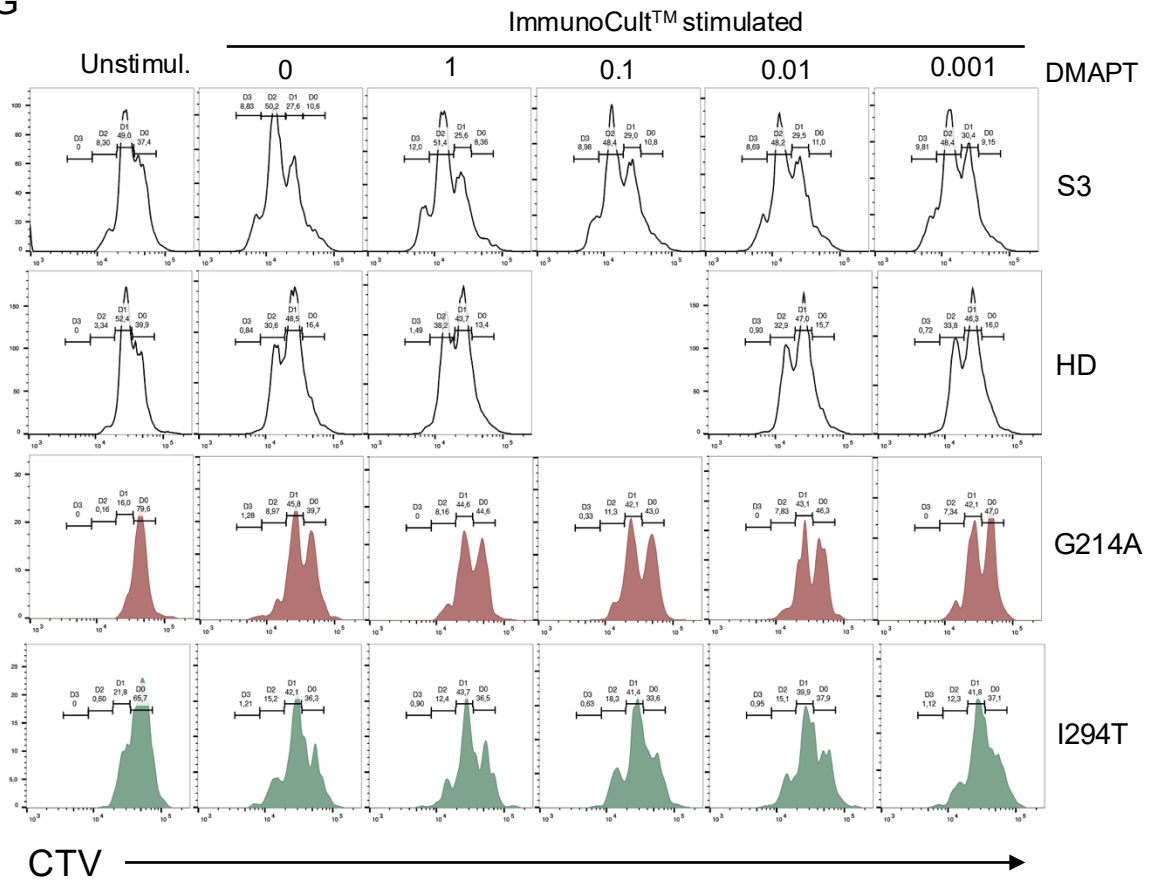

Visual Abstract

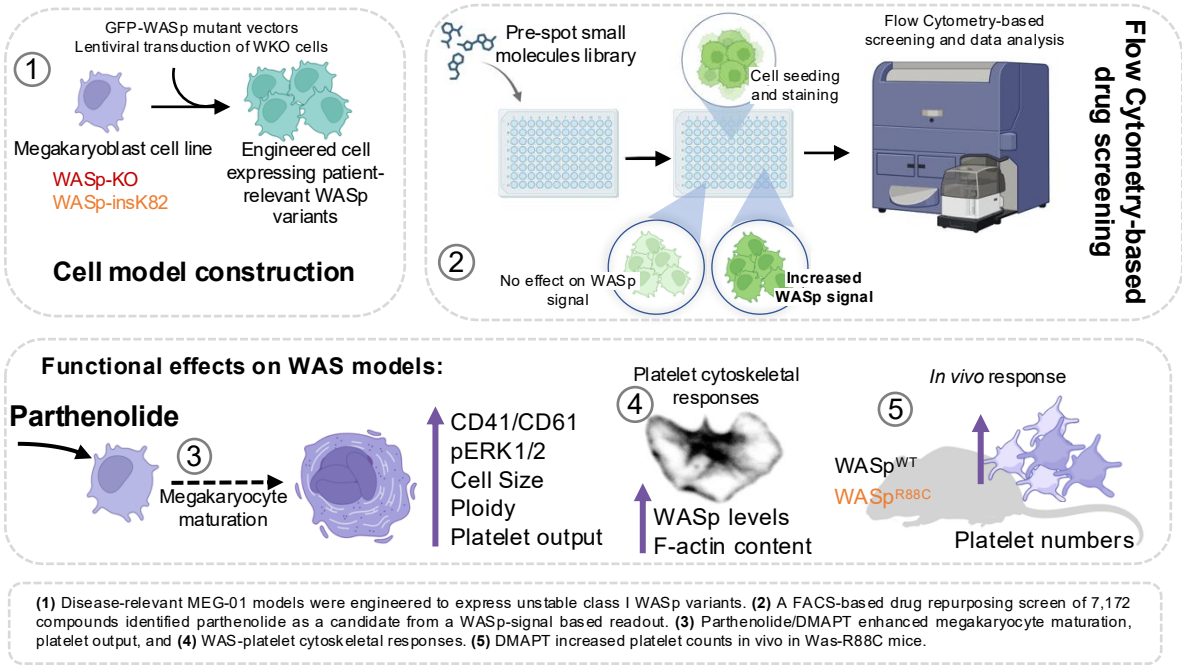
