## Supplementary material for "Parthenolide boosts megakaryocyte maturation and restores platelet responses in Wiskott–Aldrich syndrome": Methods extended

### *Human samples*

Washed platelets from healthy donors were obtained from buffy coats of anonymous blood donors processed by Karolinska Universitetssjukhuset and upon written consent. Blood from patients and family caregivers was provided upon written consent. Blood was collected in citrate tubes and received upon 24h after withdrawing. The Code of Ethics of the World Medical Association (Declaration of Helsinki) for human samples was followed; the study was approved by the institutional Ethical Committee (EPM# 2023-04773).

### *Generation of Was-R88C mice*

To generate a mouse model carrying the class I WAS-associated variant corresponding to human WASp-R86C, a knock-in point mutation was introduced into the murine *Was* locus using CRISPR-Cas9 genome editing. The murine R88C substitution corresponds to the human WASp-R86C variant. CRISPR-Cas9 targeting was performed in collaboration with Karolinska Genome Engineering and the Karolinska transgenic animal core facility. A guide RNA targeting the murine *Was* locus was designed using the CRISPRevolution sgRNA EZ Kit (Synthego). A 200-nucleotide single-stranded DNA oligonucleotide repair template (ssODN; IDT) was used to introduce the R88C mutation by homology-directed repair. The repair template also included a silent mutation generating an NdeI restriction site to facilitate genotyping. CRISPR-Cas9 components, including Cas9, sgRNA, and ssODN repair template, were delivered into C57BL/6J zygotes by electroporation. Edited embryos were transferred into pseudopregnant surrogate females. Founder mice were screened for successful knock-in by PCR-based genotyping, restriction enzyme digestion using NdeI, and confirmation by Sanger sequencing. Correctly targeted founders were bred to establish the Was-R88C mouse line. Part of this work was carried out by the SciLifeLab CRISPR Functional Genomics unit at Karolinska Institutet.

C57BL/6 wildtype, WASp knockout (WASp-KO), and WAS-R88C mice were bred under specific-pathogen-free conditions in the animal facility of KM Wallenberg, Karolinska Institutet. Mice aged 6–10 weeks were used for experimental treatments. All animal experiments performed were approved by the Stockholm North Animal Ethics Committee permit #05294-2023.

### *Cell lines*

The MEG-01 cell line (ATCC® CRL2021™) was subculture according to the ATCC data sheet.

### ***Lentiviral production and transduction***

Lentiviral vectors were produced in 293T cells using a 2<sup>nd</sup> generation lentiviral system constituted by: psPAX2, pMD2.G (AddGene), and the lentiviral transfer plasmid. Site-directed mutagenesis was performed to introduce point mutations into the transfer plasmid encoding EGFP-WASp. All plasmids were sequenced to confirm the intended modifications.

### ***Generation of WAS knockout (WKO) MEG-01 cells***

WAS KO MEG-01 cells were generated by CRISPR/Cas9. Four different gRNA sequences targeting WAS were generated and cloned into lentiGuide-Puro-P2A-EGFP-mRFPstuff (Addgene #137730). For cloning, the 25bp oligos (Table 1; ordered from Thermo Fisher Scientific) were annealed and phosphorylated with T4 PNK (NEB, 0201S) and subsequently used in a ligation reaction with Bsmbl-digested lentiGuide-Puro-P2A-EGFP\_mRFPstuf vector using Quick Ligase (NEB, M2200S). Transformed One Shot™ Stbl3 cells (Thermo Fisher, C737303) were plated on LB-ampicillin agar and incubated overnight at 37 °C. Individual colonies were inoculated into LB-ampicillin broth, grown with shaking at 37 °C for 16 h, and stored as 25% glycerol stocks. Plasmid DNA from each gRNA clone was purified using the QIAprep® Spin Miniprep Kit (Qiagen) and verified by Sanger sequencing.

MEG-01 cells were lentivirally transduced to express Cas9-BFP (Addgene #78545). Sorted BFP-positive cells were electroporated (Amaxa 4D, FF-120, homemade buffer) with lentiGuide-Puro-P2A-EGFP gRNA-cloned. After 48h, EGFP/BFP double-positive cells were sorted by flow cytometry. Mismatch cleavage assay was used for polyclonal screening to determine which gRNA that provided most detectable NHEJ (Surveyor® Mutation Detection Kit - S100, IDT). Cells targeted with gRNA1 and gRNA2 were clonally expanded. Clones with absent or low WASp levels were further sequenced to identify indels spanning 300bp upstream and downstream of the Cas9-targeted region. WASp KO cells were confirmed by WB.

### ***Libraries of molecules***

A total of 7172 compounds were screened from libraries obtained from Chemical Biology Consortium Sweden (SciLifeLab) collection. The Prestwick FDA Approved Drug Set comprehend ~1200 FDA & EMA approved drugs (current or withdrawn), with known bioavailability and safety in humans. The SPECS Drug Repurposing Set comprehend ~ 6000 Drug repurposing set (Phase 1-3, preclinical and launched) built by The Broad Institute of MIT and Harvard, Cambridge, Massachusetts, USA.

### ***Flow Cytometry-Based drug screening***

To develop the high-throughput screening assay, MEG-01 cells expressing EGFP-WASp-R86C were used. The Prestwick Chemical Library and SPECS Drug Repurposing Set molecules were pre-spotted by an automated liquid handling system. The WASp-R86C cells were seeded onto pre-spotted molecules (final concentration of 10 μM) in 96-well plates containing cycloheximide (CHX) and incubated overnight. Controls included cells treated only with CHX (negative control) and cells treated additionally with bortezomib (BTZ) (positive control). All plates included negative and positive controls to assess inter-plate consistency, ensuring a Z-factor above 0.5. Compounds resulting in an accumulation index (AI) greater than 1.2 in the primary screening were selected as positive hits. Selected positive hits underwent secondary validation using antibody-based detection of WASp (anti-WASp conjugated to mouse-AF647), with molecules tested at 3 concentrations: 5, 10 and 20μM for compounds showing low AI and 1, 5 and 10μM for the ones with high AI. Molecules that maintained consistent WASp stabilization across multiple detection methods (EGFP and antibody staining) were confirmed as validated hits.

### ***Western Blot***

Protein extracts were obtained and quantified using commercial buffers. After denaturation, 20μg protein of each sample were separated on NuPAGE™ 4–12% Bis-Tris gels (Invitrogen) in MOPS running buffer at 130 V, then transferred to PVDF membranes (Millipore) at 20 V for 90 min in Bolt™ transfer buffer with 10% methanol. Membranes were blocked 2h in 5% BSA/TBS-T and incubated overnight 4°C under shaking with anti-WASp (F8 clone, Santa Cruz, sc-365859) diluted at 1:600. After washing, membranes were incubated for 1h at room temperature with HRP-conjugated goat anti-mouse IgG diluted 1:2000. Bands were visualized using enhanced chemiluminescence (ECL Prime, GE Healthcare) and imaged on ImageQuant LAS4000. Quantification of band intensities was performed using ImageJ.

### ***Cycloheximide chase-assay***

A WASp chase assay was performed by incubating MEG-01 cells with cycloheximide (100μM, Sigma-Aldrich 239763) for different amount of time at 37°C 5% CO<sub>2</sub> incubator. Cells were collected, incubated with live/dead eFluor780 (Invitrogen, 65-0865-14), fixed with PFA 4% and intracellularly stained with primary anti-WASp antibody (F8 clone, Santa Cruz, sc-365859) followed by staining with secondary antibody goat anti-Mouse IgG (H+L) secondary antibody, Alexa Fluor 647 (Invitrogen).

### ***Immunofluorescence***

Sterile coverslips were coated with 100 µg/mL plasminogen-depleted human fibrinogen (Merck, 341578) for 2h at room temperature or overnight at 4°C, then washed twice with Milli-Q water and blocked for 1h in 1% BSA. MEG-01 cells, pre-treated ± PTW08, were seeded onto the coated coverslips and incubated overnight at 37°C, 5% CO<sub>2</sub>. Coverslips were fixed in 4% PFA prepared in PEM buffer (8 mM PIPES pH 6.8, 1 mM EGTA, 1 mM MgCl<sub>2</sub>) for 20 min at room temperature, after blocking and permeabilization. Nuclear and F-actin staining was performed with DAPI and Alexa 568-Phalloidin in TBS for 20 min, followed by 3 TBS washes. Coverslips were mounted on ProLong™ Glass Antifade (Invitrogen, P36984) and cured overnight, sealed with nail polish, and stored at 4 °C. Confocal z-stacks (0.5 µm steps) were acquired with a Zeiss LSM 880 using a 63× oil-immersion objective, imaging at least 30 cells per condition.

#### ***Platelet activation assay***

Briefly, PRP was prepared by centrifuging the blood at 200 × g for 20 minutes at room temperature, without brake. Two-thirds of the upper layer were collected and diluted with citrate buffer containing PGE1 (1 µg/mL, Sigma Aldrich), then centrifuged at 800g for 10 minutes. The platelet pellet was gently resuspended in Tyrode's buffer with 0.3% BSA. Platelets (5x10<sup>6</sup>) were incubated with mouse anti-human-CD41 Alexa Fluor 700 (HIP8 clone; BioLegend) at 1:100 for 10 minutes and then stimulated with 1 U/mL thrombin (Sigma Aldrich) and fixed at the appropriate time with Tyrode's buffer + PFA 4% for 10 minutes. Fixed cells were permeabilized and stained intracellularly for WASp and F-actin analysis. Samples were acquired on BD X-20 LSR Fortessa.

#### ***Megakaryocyte Ploidy Analysis***

Cells were prepared at 1 × 10<sup>6</sup> cells per sample and, for some samples, stained for extracellular markers and viability prior to fixation. Pelleted cells were fixed in PFA 4% for 20 min RT, then washed. The pellet was resuspended in 300 µL DAPI solution (20 µg/mL in 0.1% Triton X-100/PBS) and incubated in the dark for 30 min at room temperature. Samples were kept cold and protected from light, without further washes, until the acquisition. Data was acquired on BD X-20 LSR Fortessa, set to linear scale under low pressure flow. DNA content histograms were analyzed to determine cell-cycle distribution.

#### ***Mean ploidy-index***

DNA ploidy was assessed by flow cytometry in gated megakaryocytes. Mean ploidy index was calculated within the polyploid megakaryocyte population, including 4N, 8N, 16N, 32N, and

64N fractions. The mean ploidy index was calculated as the weighted average of each ploidy class using the following formula:

Mean ploidy index = $\sum (P_i \times N_i)/100$ , where $P_i$ represents the percentage of cells in each ploidy class and $N_i$ represents the corresponding ploidy value (2N, 4N, 8N, 16N, 32N, or 64N).

### ***Primary Megakaryocytes culture***

Bone marrow cells were isolated from femurs and tibias of WT, WASp-R88C, and WASp-KO mice. Hematopoietic stem and progenitor cells were enriched from total bone marrow using EasySep™ Mouse Hematopoietic Progenitor Cell Isolation Kit according to the manufacturer's instructions. Isolated cells were cultured under conditions adapted from the mouse HSC expansion protocol as described (25). Cells were first expanded for 5–6 days in expansion medium supplemented with recombinant murine stem cell factor (SCF, 50 ng/mL) and thrombopoietin (TPO, 100 ng/mL). After expansion, cells were differentiated toward the megakaryocyte lineage for an additional 7–9 days in medium supplemented with TPO alone (100 ng/mL). Cultures were maintained at 37°C with 5% CO<sub>2</sub>, and fresh cytokine-containing medium was added or replaced as required. Mature megakaryocytes were analyzed by flow cytometry based on expression of CD41, CD61, and CD42d, DNA ploidy, and platelet production in the culture supernatant. For drug-treatment experiments, parthenolide or DMAPT was added during the megakaryocyte differentiation phase at the indicated concentrations. Vehicle-treated cultures were included as controls.
